# Bromodomain-containing Protein 4 Regulates Innate Inflammation in Airway Epithelial Cells via Modulation of Alternative Splicing

**DOI:** 10.1101/2023.01.17.524257

**Authors:** Morgan Mann, Yao Fu, Xiaofang Xu, David S. Roberts, Yi Li, Jia Zhou, Ying Ge, Allan R. Brasier

## Abstract

Bromodomain-containing Protein 4 (BRD4) is a transcriptional regulator which coordinates gene expression programs controlling cancer biology, inflammation, and fibrosis. In airway viral infection, non-toxic BRD4-specific inhibitors (BRD4i) block the release of pro-inflammatory cytokines and prevent downstream remodeling. Although the chromatin modifying functions of BRD4 in inducible gene expression have been extensively investigated, its roles in post-transcriptional regulation are not as well understood. Based on its interaction with the transcriptional elongation complex and spliceosome, we hypothesize that BRD4 is a functional regulator of mRNA processing. To address this question, we combine data-independent analysis - parallel accumulation-serial fragmentation (diaPASEF) with RNA-sequencing to achieve deep and integrated coverage of the proteomic and transcriptomic landscapes of human small airway epithelial cells exposed to viral challenge and treated with BRD4i. The transcript-level data was further interrogated for alternative splicing analysis, and the resulting data sets were correlated to identify pathways subject to post-transcriptional regulation. We discover that BRD4 regulates alternative splicing of key genes, including Interferon-related Developmental Regulator 1 (*IFRD1*) and X-Box Binding Protein 1 (*XBP1*), related to the innate immune response and the unfolded protein response, respectively. These findings extend the transcriptional elongation-facilitating actions of BRD4 in control of post-transcriptional RNA processing in innate signaling.

## 1. Introduction

Bromodomain-containing protein 4 (BRD4) is a key regulator of development, tumorigenesis, inflammation, and fibrosis in mammalian organisms [1–5]. In the context of airway inflammation, aeroallergens and airway pathogens trigger NF-*κ*B/BRD4-mediated innate inflammation via activation of upstream pattern recognition receptors [6]. These include the toll-like receptors (TLRs) and the RIG-Helicase (RIGI), which induce the proteolytic degradation of the NF-*κ*B inhibitory subunit (NF*κ*BIA), resulting in cytoplasmic release and nuclear translocation of NF-*κ*B subunit p65 (RelA), where it interacts with BRD4 and the latent positive transcription elongation factor (pTEFb) [7]. Upon BRD4 binding, pTEFb dissociates from inhibitory 7SK RNA/HEXIM, and the resulting complex phosphorylates promoter-proximal paused RNA polymerases and negative elongation factors. In this way, PTEFb produces rapid transcriptional elongation of fully spliced, immediate-early NF-*κ*B-dependent genes, resulting in rapid and efficient activation of pro-inflammatory cytokines, chemokines, and anti-viral interferons. This process provides a key initial barrier to viral infection, and recruits a robust cellular immune response for prolonged protection by stimulating adaptive immunity. Additionally, prolonged BRD4 activation results in epithelial plasticity, basement membrane remodeling, and fibrosis [8].

BRD4 derives its name and primary function from its tandem, acetyl-lysine binding bromodomains (BDs), which facilitate interactions with acetylated histones and other acetylated proteins [6, 9, 10]. This characteristic enables BRD4 to bind tightly to transcriptionally active DNA, serving as an epigenetic ”bookmark” [11]. Protein interaction studies have observed that BRD4 binds not only to acetylated NF-*κ*B p65/RelA, but forms a dynamic multi-component complex with the AP-1 transcription factors, members of the Mediator complex, SWI-SNF chromatin-remodeling complex, and RNA-splicing factors [12, 13]. Understanding the BRD4 protein-protein interaction network provides novel insights into the pleiotropic functions of the BRD4 complex.

The development of potent and selective small-molecule inhibitors of the BRD4 BDs has enabled systematic interrogation of dynamic BRD4-acetylated protein interactions [12]. Interestingly, BRD4 interactions with members of the spliceosome complex can be disrupted by inhibitors of the acetyllysine binding pocket. The spliceosome is a multi-megadalton complex of 5 small nuclear RNAs (snRNAs) and approximately 100 proteins that function together to process messenger RNA precursors (pre-mRNA) and process them into mature mRNA [14]. This involves catalytic removal of non-protein coding intronic regions of the pre-mRNA, and re-ligation into a contiguous, translatable mRNA. Key protein subunits of the spliceosome include the small nuclear ribonucleoproteins (snRNPs) U1 and U2, which form a pre-spliceosome complex at the sites of intronic regions, and the U4, U5, and U6 snRNPs which complete the spliceosome and initiate splicing through a U2/U6-dependent mechanism [15, 16]. Numerous additional snRNPs and splicing factors contribute structural and functional roles to the complex, coupling mRNA splicing to processes such as transcription and mRNA export [17, 18]. Alternative splicing of exon regions (a.k.a. exon-skipping) is also possible, and gives rise to significant diversity in the transcriptome [19]. Importantly, the role of BRD4 in alternative splicing is not fully understood.

Respiratory Syncyctial Virus (RSV) is an enveloped, single-stranded, negative-sense RNA virus that is the largest cause of pediatric hospitalizations worldwide [20, 21]. RSV thus represents a significant burden on healthcare systems, especially as cases have surged in the wake of the SARS-CoV-2 pandemic [22]. RSV infects the epithelial cells of the respiratory tract, and induces rapid activation of the NFκB-BRD4-PTEFb pathway [2, 6, 12, 23–27]. Previous single molecule RNA sequencing studies have discovered that RSV replication induces substantial alternative splicing events, including intron retention and alternative polyadenylation site utilization [28]. However, the mechanisms by which RSV induces alternative splicing are not known.

In this study, we investigate BRD4-mediated alternative splicing in the context of RSV-induced innate inflammation, using a multi-omics approach combining in-depth short-read RNA-sequencing with high-throughput trapped ion mobility (TIMS) mass spectrometry. This combined approach enables a deep profiling of both the transcript- and protein-level abundance changes that result from alternative splicing events. Our results demonstrate that BRD4 regulates RSV-induced intron retention, consequently modulating the abundance of associated co-regulators of innate inflammation. Furthermore, we demonstrate that these changes result in decreased activation of NF*κ*B-mediated gene expression, independent of BRD4 inhibition.

## 2. Results

### 2.1. BRD4 inhibition alters transcriptome-wide intron-retention and ORF length in airway epithelial cells

To investigate the effects of BRD4 inhibition and viral infection on alternative splicing in the airway epithelium, we analyzed data from *Xu et. al.* [28] to discover alternative splicing events. This data was collected in mock or RSV-infected telomerase-immortalized human airway epithelial cells (hSAECs) in the absence or presence of treatment with the specific BRD4 inhibitor ZL0454. hSAECs are a well-established airway epithelial model to characterize the mechanisms of innate airway inflammation. These cells maintain stable epithelial morphology in monoculture, and reproduce both genomic and proteomic signatures of primary cells without early senescence [25]. ZL0454 is a potent competitive BD inhibitor highly selective for BRD4 over other members of the bromodomain family [29, 30].

This alternative spliceform analysis revealed 2001 genes and over 3635 unique transcripts with significant isoform switching events in at least one comparison. 1008 genes had induced isoform switching due to RSV infection, and 822 genes were associated with isoform switching due to BRD4 inhibition in RSV-infected cells (Figure 1). BRD4i-associated splicing events were subjected to PANTHER gene ontology analysis [31, 32], indicating significant changes to kinetochore assembly and nucleic-acid/base-pair metabolism. 314 genes overlapped between conditions, suggesting switch events regulated by both RSV and BRD4, however, no PANTHER ontology terms were significantly enriched among these genes. We conclude that BRD4 does not alter alternative splicing in a pathway-concerted manner. Splicing events were further analyzed by association with transcript-level features. RSV-induced splicing events were associated with predominantly shorter Open Reading Frames (ORFs) and noncoding transcripts. In contrast, BRD4 inhibition reversed this pattern, and led to predominantly longer ORFs and retention of intronic regions. A full list of detected transcripts and splicing events is available in Table S1.

**Figure 1.**
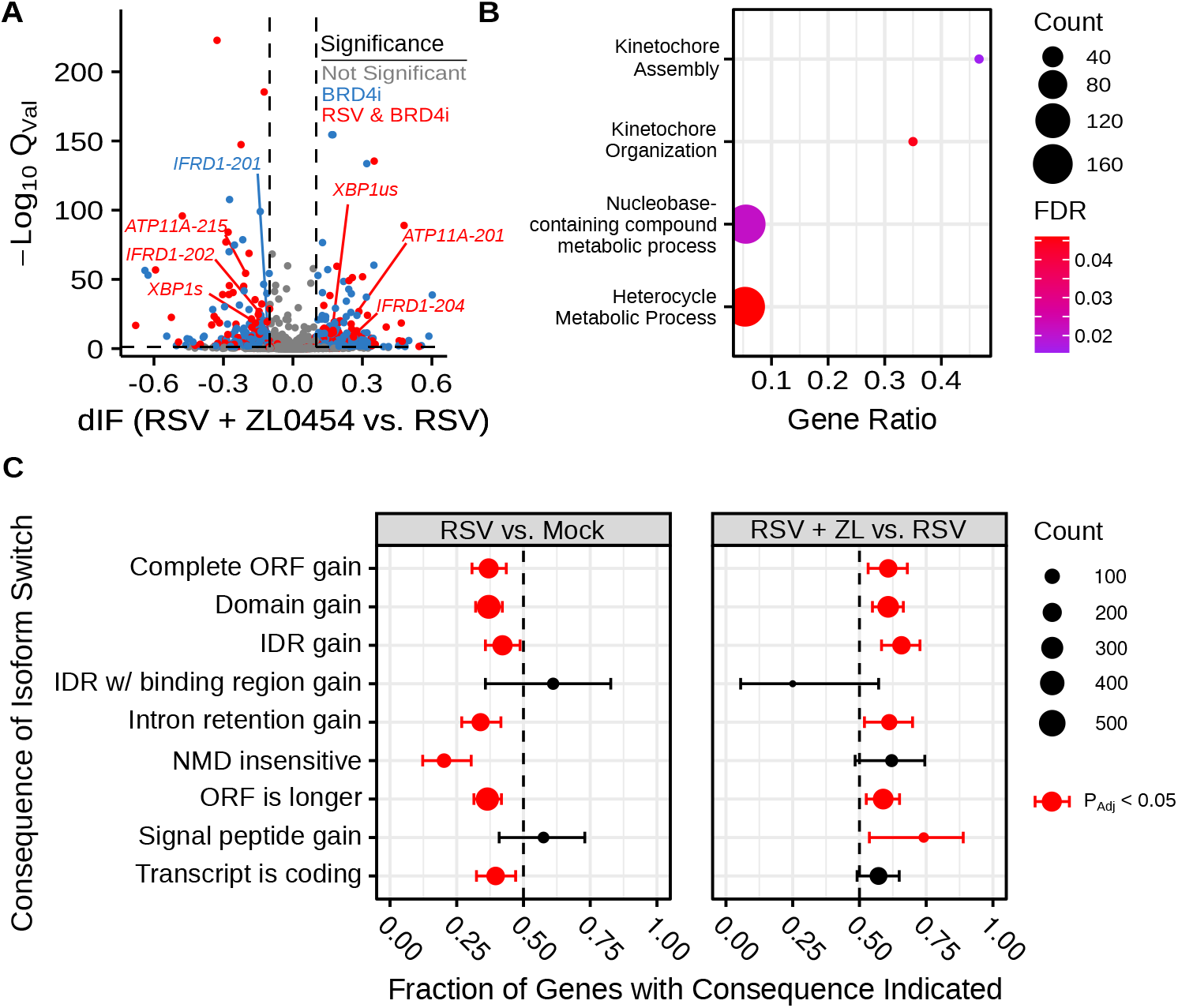
BRD4 inhibition alters alternative splicing in the context of airway viral infection. (A) Volcano plot indicating differential isoform fraction (dIF) and isoform switch q-value of isoform switching events with predicted gene-level consequences. Non-significant changes (*Q_Val_* > .05 or |dIF| < 0.1 are plotted in grey. Significant changed transcripts also found to be significantly changed due to RSV-infection are plotted in red. Significant switch events only detected due to BRD4 inhibition are plotted in blue. (B) Gene Ontology Biological Process Over-representation analysis of BRD4i-mediated splicing events. Count refers to the number of alternatively spliced genes detected in the significantly over-represented pathway. (C) Transcript-level consequences of alternatively spliced genes as induced by RSV or BRD4 inhibition.

### 2.2. BRD4 regulates alternative splicing of key innate inflammation coregulators

Manual evaluation of this dataset revealed that BRD4i was associated with sequence retention in several genes of note: X-box Binding Protein 1 (*XBP1*) and Interferon-related Developmental Regulator 1 (*IFRD1*) (Figure 2). The *XBP1* gene codes for a key transcription factor in the unfolded protein response [33], but must first be spliced to an active form, *XBP1s,* via the excision of a 26 nucleotide sequence [34]. This results in a frame shift that allows the mRNA to be translated into the functional transcriptional regulator. XBP1 has been shown to activate expression of the Interleukin-6 (IL6) cytokine [35, 36], a key cytokine in the innate immune response. In our dataset, we observe that BRD4i blocks this splicing event, maintaining *XBP1* mRNA in its longer, inactive form (Figure 2). This coincides with an overall reduction in *XBP1* gene expression to dramatically reduce the abundance of *XBP1s* transcripts. We validate this result using q-RT-PCR primers targeting exon 4 of *XBP1s* (Figure 3C).

**Figure 2.**
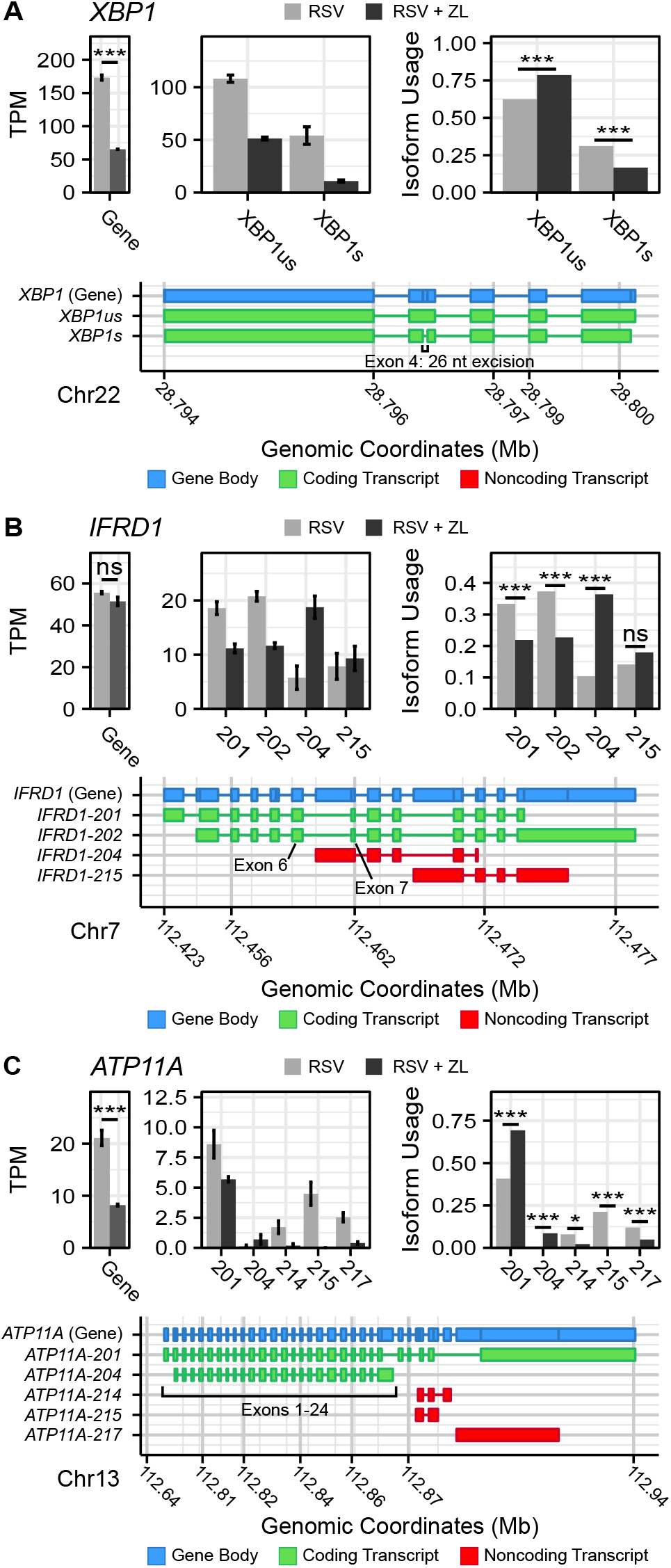
BRD4 inhibition induces isoform switching in key innate inflammation co-regulators. Transcript maps, RNA-seq quantitation in Transcripts-per-Kilobase Million (TPM), and isoform usage of detected transcripts belonging to (A) *XBP1,* (B) *IFRD1,* and (C) *ATP11A.* Mapped transcripts are plotted in compressed genomic coordinates and color coded according to gene body and coding status. Barplots represents the mean of n=4 replicates per group. Error bars indicate the 95% confidence interval. Transcripts with less than 5% of total detected transcript abundance were omitted from this figure. \**P_Adj_* < 0.05; \*\*\**P_Adj_* < 0.001.

**Figure 3.**
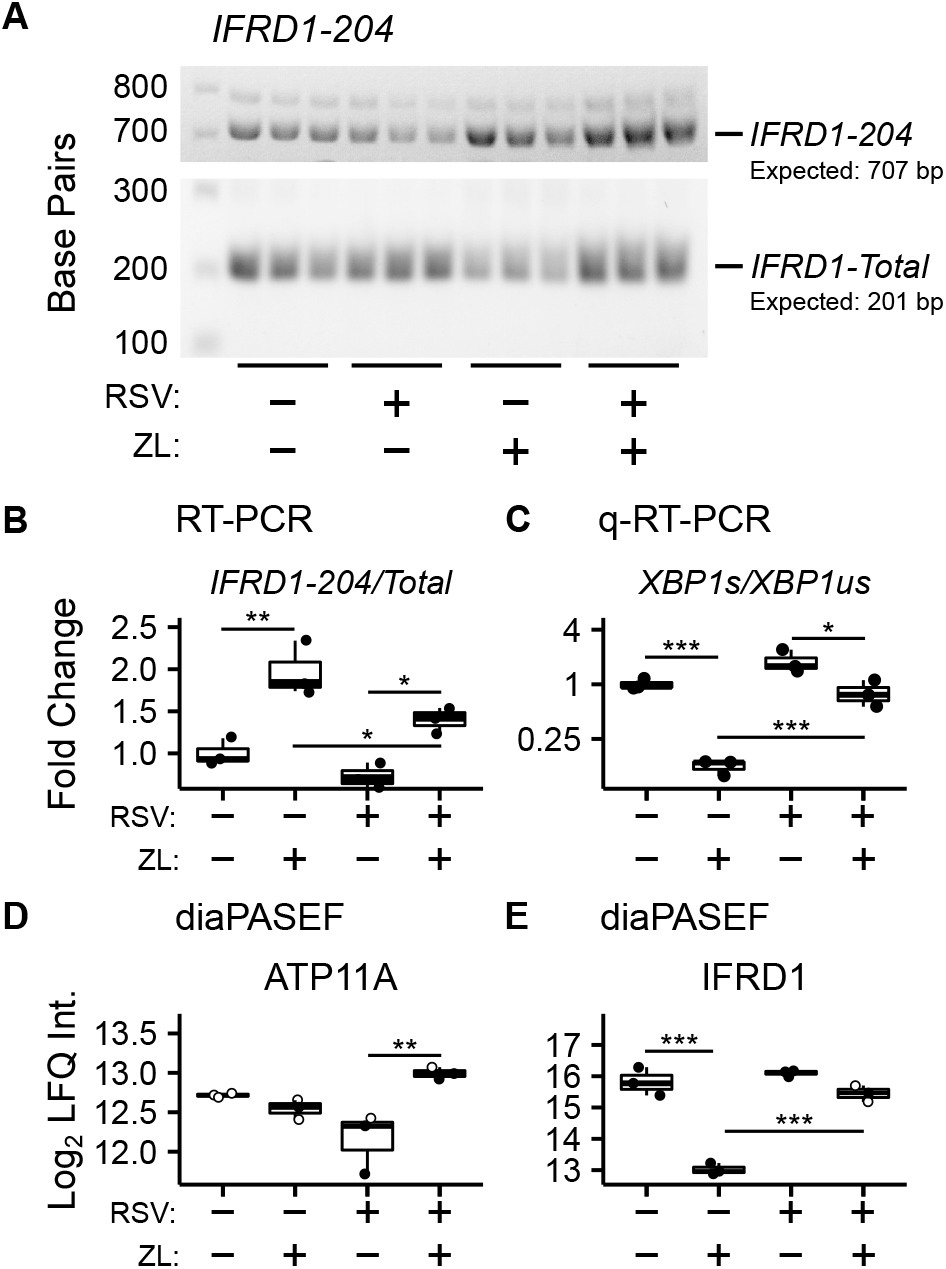
Validation of BRD4i-induced alternative splicing. (A) RT-PCR Amplification of IFRD1-204 and IFRD1-Total. (B) Gel Densitometry quantitation of IFRD1-204/IFRD1-Total Ratio. (C) q-RT-PCR quantitation of XBP1s/XBP1-Total Ratio. (D) Protein-level (diaPASEF) quantitation of ATP11A and IFRD1 abundance. Boxplots represent median and interquartile ranges of n=3 biological replicates per experimental group. Data points imputed during protein differential abundance analysis are rendered in grey. *\*P_Adj_* < 0.05; *\*\*P_Adj_* < 0.01; \*\*\**P_Adj_* < 0.001.

*IFRD1*, in contrast, is a poorly-studied, histone deacetylase-associated transcriptional regulator. Specifically, its function has been linked to the acetylation of NF-*κ*B p65/RelA [37, 38] - a key activator of the innate response, which interacts with BRD4 when acetylated at lysine-310. In our model, we observe that BRD4i does not change overall gene expression of *IFRD1*. However, BRD4i does result in an alternative transcription start site located in the intron between exons 6 and 7. This excises exons 1-6, and results in a non-functional, non-coding transcript (Figure 2A). This result was confirmed by RT-PCR, using primers which span the intronic region and the canonical exons (Figure 3AB).

In addition, we observe splicing events to the membrane-bound ATPase Phospholipid Transporting 11A (*ATP11A),* a member of the P4-ATPase Flippase complex, which is responsible for transporting a variety of phospholipids across the plasma membrane [39]. Importantly, this complex has been linked to innate immunity via regulation of toll-like-receptor (TLR) re-uptake into the plasma membrane from endocytic vesicles [40]. Curiously, we observe that while BRD4i reduces the gene-level abundance of *ATP11A,* it predominantly does so by ablating non-coding transcript variants. Accordingly, the abundance of coding transcripts remains predominantly unchanged (Figure 2A). Due to the extreme heterogeneity in non-coding *ATP11A* transcripts, no RT-PCR assay could be designed to validate this event.

### 2.3. Protein-level validation of alternative splicing

To determine corresponding protein-level changes, we analyzed the proteome of hSAECs under identical biological conditions. Cells were pre-treated with ZL0454 or DMSO for 18 hours prior to infection with RSV at a multiplicity of infection (MOI) of 1. Control cells were left uninfected and untreated with ZL0454. 24 hours later, cells were harvested and processed for shotgun proteomic analysis by Data Independent Analysis - Parallel Acquisition Serial Fragmentation (diaPASEF) [41]. This enabled quantitation of ~8300 proteins, including 6 RSV proteins (1B, 1C, F, G, L, and P), with ~94.6% data completeness prior to filtering (Figure S1AD). Across 5 contrasts (RSV vs. Control; ZL0454 vs. Control; RSV + ZL0454 vs. Control; RSV + ZL0454 vs. RSV; RSV + ZL0454 vs. ZL0454), we observed 9332 statistically significant protein changes. Comparison of RSV-treated cells with ZL0454-treated cells was excluded from the differential abundance analysis to preserve statistical power. Reproducibility of biological replicates was high, as the lowest intra-group pearson correlation score was 0.938, and the median coefficient of variation ranged from ~1-2% within groups (Figure S1BC). A full breakdown of differentially abundant proteins by contrast is available in Table S2. ~7500 genes had matched protein-level and transcript-level measurements. IFRD1 and ATP11A were among the proteins that were differentially abundant based on treatment with the BRD4 inhibitor (Figure 3D). XBP1/XBP1s was not detected by the proteomic analysis, most likely due to the extremely low abundance of this transcription factor.

In this analysis, we observed that the abundance of IFRD1 protein remains unchanged due to RSV-infection, but drops 8-fold when treated with the ZL0454 alone. IFRD1 protein abundance is also slightly reduced by the inhibitor under the RSV condition, however, the abundance could not be reliably assayed, as 2 of 3 values were missing in the proteomic dataset and were imputed for differential analysis (Table S3). This result is consistent with the prediction from the alternative splicing analysis, in which BRD4i transitions the transcripts to a non-protein coding variant. ATP11A protein abundance, as predicted, remains unchanged when treated with the BRD4 inhibitor alone. Interestingly, however, ATP11A abundance actually increases in abundance (~75%) in the combined combined RSV + ZL0454 condition relative to RSV-treatment alone, despite a ~2.5-fold decrease at the total RNA level, and reverses a non-significant trend towards reduced ATP11A protein observed in RSV-treated cells. As the total abundance of the functional ATP11A transcript did not increase, this suggests that ATP11A is regulated by additional, post-translational mechanisms of gene regulation.

### 2.4. IFRD1-depletion reduces basal innate inflammation

As IFRD1 function has been linked directly to RelA acetylation [37, 38], and RelA Lysine-310 acetylation (K310Ac) is required for BRD4-mediated inflammation, we were interested in assaying the effects of IFRD1 depletion on the process of innate activation in airway epithelial cells. Pooled siRNA targeting IFRD1 was transfected into hSAECs, and peak knockdown efficiency was determined by western blot to occur after 72 hours (**Figure S2**).

IFRD1-depleted hSAECs were then stimulated with the Toll-like Receptor 3 (TLR3)-ligand, polyinosinic-polycytidilyic acid (polyIC), a synthetic viral replication intermediate that activates innate inflammation [42, 43]. TLR3 signaling proceeds through BRD4:RelA-mediated transcriptional elongation, and accordingly, *IL6* was chosen as a marker of innate activation. In this knockdown model, we observed that expression of *IL6* was reduced ~4-fold during early innate activation. (Figure 4). However, this effect was no longer significant by 60 minutes post-stimulation, suggesting activation of compensatory pathways.

**Figure 4.**
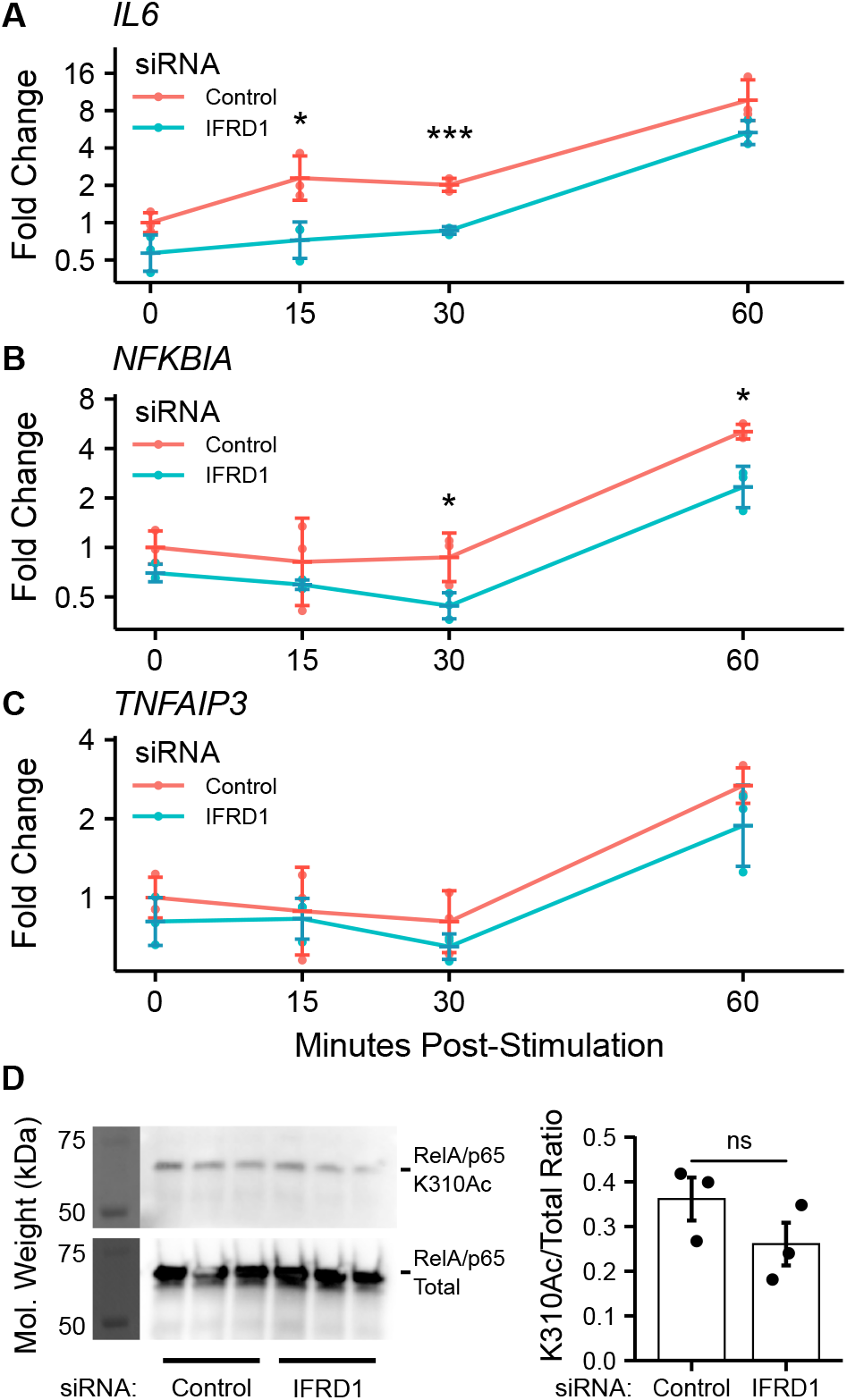
IFRD1 knockdown reduces NF-*κ*B-mediated expression in hSAECs. Timecourse q-RT-PCR experiment measuring (A) IL6, (B) *NFK-BIA,* and (C) *TNFAIP3* gene expression in hSAECs depleted of IFRD1. Data represents the geometric mean fold change ± standard error of n=3 biological replicates per experimental group. *\*P_Adj_* < 0.05; *\*\*\*P_Adj_* < 0.001. (D) Western blot demonstrating persistent IFRD1 knockdown throughout the timecourse experiment.

We also monitored the gene expression of the TNF*α*-induced Protein 3 *TNFAIP3* and NF-*κ*B Inhibitor Alpha *NFKBIA* genes, which encode the A20 and IK*βα* proteins and are both NF-*κ*B-dependent and highly-induced by innate activation [44]. *NFKBIA* expression was similarly reduced by IFRD1 knockdown, but the effect did not ablate by 60 minutes poststimulation; *TNFAIP3* displayed only a non-significant trend towards reduced gene expression. IFRD1 knockdown remained stable throughout the duration of the experiment. Curiously, the molecular weight of IFRD1 was observed at ~75 kilodaltons (kD), which is ~25 kD larger than the predicted from the amino acid sequence, potentially indicating the presence of un-annotated post-translational modifications.

Curious as to whether the inhibitory effects of IFRD1 knockdown behavior were a consequence of altered RelA-Acetylation, we immunoprecipitated RelA from A549 alveolar epithelial cells and attempted to measurethe K310Ac mark via immuno-blotting. A549 cells have been extensively utilized for analysis of airway innate responses and maintain characteristics of type II alveolar cells [45, 46]. To our surprise, we found that the abundance of the RelA K310Ac mark was not significantly effected by knockdown of IFRD1. This result is inconsistent with previous works conducted in keritinocytes, in which RelA K310 acetylation was increased in response to IFRD1 knockdown [38], and suggests that IFRD1 has cell-type or context-specific mechanisms of innate inflammatory regulation.

### 2.5. BRD4 regulates the abundance of spliceosome components in airway epithelial cells

In our analysis of alternative splicing, we sought to determine mechanisms for these changes, and accordingly examined the effects of BRD4i on the abundances of spliceosome components and splicing factors. Consistent with previous reports [13, 47], the data from *Xu et. al.* [28] did not indicate major, BRD4-mediated changes to the abundance of splicing factors. However, a 2-Dimensional Annotation analysis [48] of RNA-sequencing data with protein-level quantitation indicated significant disagreement between protein- and transcript-level abundances in proteins related to the spliceosome (Figure 5A). In general, spliceosome-related transcripts marginally increased in abundance after BRD4-inhibition, but matching protein abundance decreased by a much larger magnitude.

**Figure 5.**
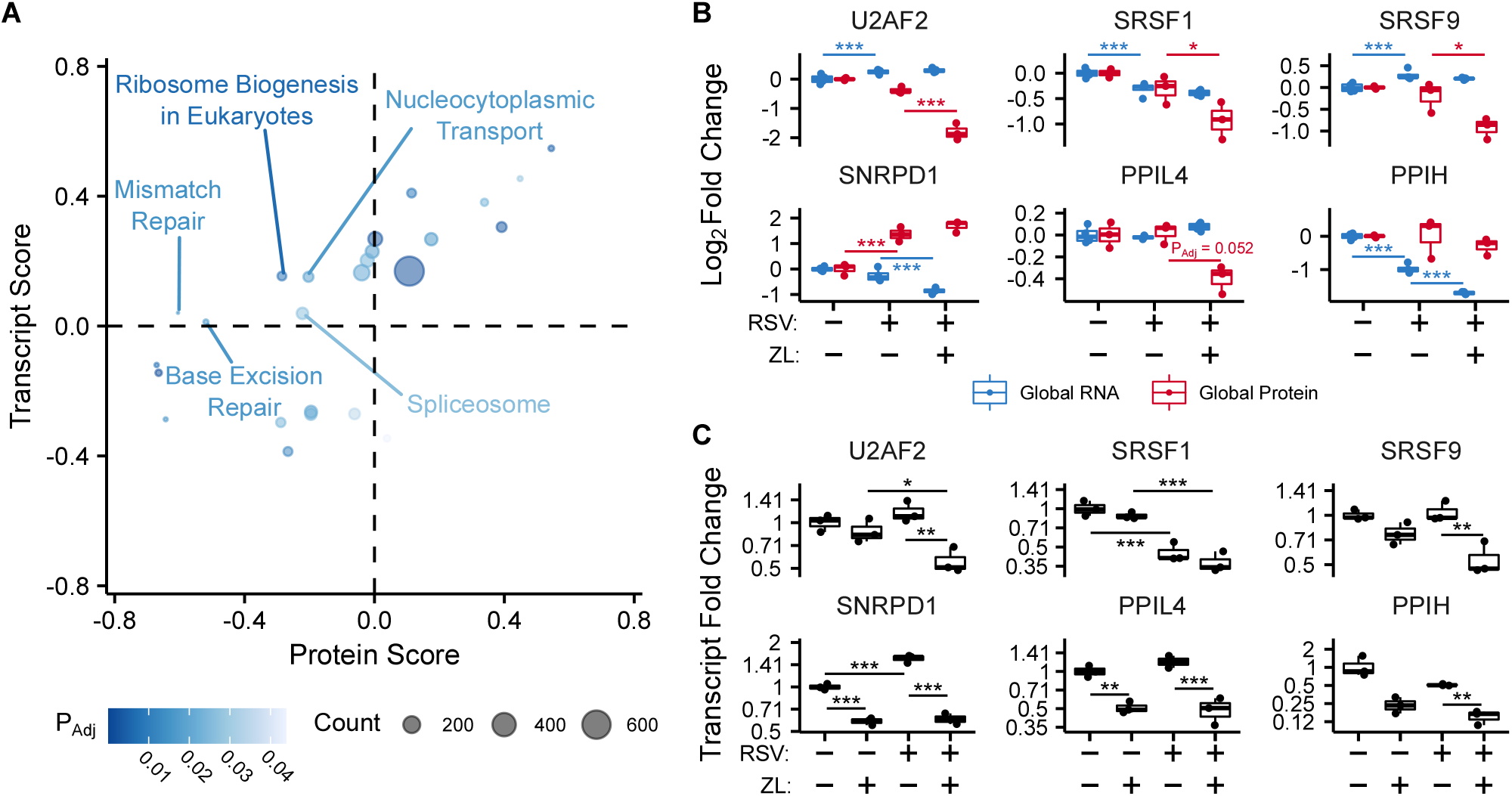
BRD4 regulates the transcript- and protein-level abundance of core splicing factors. (A) 2-dimensional annotation analysis of matched RNA-sequencing and proteomic data. Significant (FDR < 0.05) annotation terms are plotted on the graph. Transcript/Protein Scores are proportional to relative abundance. (B) Protein and transcript-level measurements of core splicing factors, as quantified by diaPASEF and RNA-sequencing. The ZL0454-only condition has been omitted as it was not included in *Xu et. al.* [28]. Boxplots represent median and interquartile range. Protein data represents n=3, and RNA-sequencing represents n=4 biological replicates per experimental group. (C) q-RT-PCR validation of transcript-level splicing factor abundances. Boxplots represent median and interquartile range of n=3 biological replicates per experimental group. *\*P_Adj_* < 0.05; *\*\*P_Adj_* < 0.01; *\*\*\*P_Adj_* < 0.001.

As BRD4 is known to interact with the spliceosome [12, 13], this data initially suggested that BRD4 regulates the abundance of spliceosome components through post-transcriptional mechanisms. However, q-RT-PCR validation contradicted the RNA-sequencing results in 4 of 6 examined genes, including the core splicing factor U2AF subunit 2 (U2AF2), as well as several serine-responsive splicing factors (SRSF1/SRSF9) and Peptidylprolyl Isomerase Like 4 (PPIL4) (Figure 5C). Interestingly, each of these cases was associated with reduced gene-product abundance. In contrast, q-RT-PCR quantitation for Small Nuclear Ribonucleoprotein D1 Polypeptide (SNRPD1) and Peptidylprolyl Isomerase H (PPIH) demonstrated decreased transcript abundance, as predicted by the RNA-sequencing results, despite unchanged protein-level abundance. This may indicate that SNRDP1 and PPIH proteins are stable during the time-frame of the conducted experiments, however, the discrepancy in transcript-level metrics for *U2AF2* and other splicing factors can not be easily explained. Given the general agreement with other sequencing studies [13, 47], we suggest that this behavior could result from the presence of unannotated transcript variants which can not be quantified by quasi-mapping alignment tools [49], from sequencing artifacts introduced during sample preparation [50], or from violation of standard differential gene expression assumptions [51]. We conclude that BRD4 transcriptionally controls the abundance of core spliceosome components in human small airway epithelial cells.

### 2.6. BRD4 regulates XBP1s splicing through transcriptional control of IRE1*α*

In addition to spliceosome-associated splicing factors, we were also able to quantify the effects of BRD4i on the endoplasmic reticulum (ER)-bound splicing factor Inosital-requiring Enzyme 1 *α* (IRE1*α*). IRE1*α* is an atypical protein kinase and endoribonuclease which is seated in the membrane of the ER, and uniquely splices *XBP1* transcripts into *XBP1s*. Interestingly, we observe that BRD4 inhibition blocks a ~75% increase in the abundance of IRE1*α* due to RSV infection, which coincides with a reduction to *IRE1α* gene expression in the same contrast (Figure 6). The effect of ZL0454 alone was not measured in *Xu et. al.* [28] and was not statistically significant at the protein-level. However, we note that 2 of 3 biological replicates in this condition were imputed during differential abundance analysis; accordingly, the abundance could not be reliably assayed.

**Figure 6.**
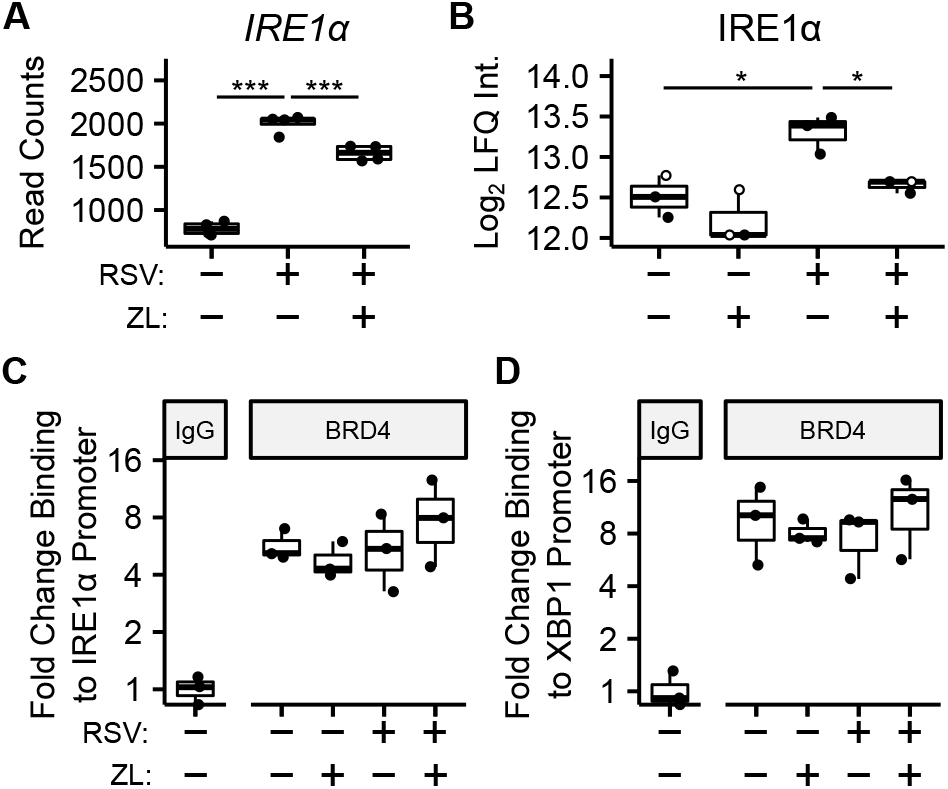
BRD4 controls the transcript- and protein-level abundance of IRE1*α*. (A) RNA-sequencing read counts of summarized *IRE1a* transcripts. (B) Protein-level (diaPASEF) quantitation of IRE1*α* protein. Data points imputed during protein differential abundance analysis are rendered in grey. (C) ChIP-PCR analysis of BRD4 binding to the *IRE1a* promoter. (D) ChIP-PCR analysis of BRD4 binding to the *XBP1* promoter. Boxplots represent median and interquartile ranges of n=4 (RNA-seq) or n=3 (diaPASEF, ChIP) biological replicates per experimental group. *\*P_Adj_* < 0.05; *\*\*\*P_Adj_* < 0.001.

Given that BRD4 inhibition reduces both *XBP1* and *IRE1α* gene expression, these results suggested that BRD4 controls *XBP1* splicing through transcriptional control of *IRE1α*. To further investigate this hypothesis, we conducted chromatin immunoprecipitation (ChIP) assays to measure the abundance of BRD4 protein on the promoters of both genes in hSAECs infected with RSV and treated with ZL0454. This demonstrated a 4-8–fold enrichment of BRD4 on the promoters of these genes, but to our surprise, neither RSV-infection nor BRD4i resulted in any perturbations to its presence. This constitutive enrichment suggests that BRD4 plays a secondary role in activating *IRE1α* and *XBP1* expression, perhaps through stabilization of additional transcriptional regulators at the promoters of these genes.

## 3. Discussion

Respiratory Syncytial Virus (RSV) is a ubiquitous human pathogen for which no effective vaccine is approved, and is the single largest cause of pediatric hospitalizations in the United States [21]. Replication of RSV or other airway viruses in the lungs induce airway epithelial cells to mount a robust innate inflammatory response to reduce viral replication and recruit cellular immune cells to the site of infection. This process represents a significant anti-viral barrier, and plays a major role in the resolution of the disease. The innate inflammatory pathway begins with pattern-recognition receptors (PRRs), such as Toll-like receptors on the cell surface and the retinoic-acid inducible gene (RIG-I) in the cytosol [6, 24], and largely proceeds through NF-*κ*B and IRF transcription factors [27, 52, 53]. These pathways converge on BRD4, which is an epigenetic scaffold that interacts with elements of the positive transcriptional elongation factor (pTEFb) [27]. In this manner, BRD4 enables the transcriptional elongation and rapid expression of interferons, cytokines, and other genes with polymerases paused on their promoters [54]. However, as innate inflammation is highly pleiotropic and stimulus-specific, and the exact mechanisms by which BRD4 facilitates transcription in a gene- and time-resolved manner have yet to be elucidated. BRD4 has also been shown to interact with components of the spliceosome and ribonuclear protein complexes [12, 13], suggesting a role in post-transcriptional regulation of gene products. These interactions have also been shown to be sensitive to BRD4 inhibition. In this work, we further demonstrate that BRD4 inhibition alters the splicing landscape of airway epithelial cells, resulting in altered abundances of proteins associated with the innate immune response and innate inflammation. These alternative splice events modulate innate signaling through the TLR3-NFκB signaling pathway and the unfolded protein response. Our data provide novel insights into the diverse anti-inflammatory actions of BRD4 inhibition.

### 3.1. Pleiotropic activities of BRD4 in transcription-coupled RNA processing

Control of innate-inducible RNA Pol II-dependent gene expression is coordinated through a series of integrated steps including promoter initiation, elongation, RNA processing, termination and export [55]. Although virtually all of these steps are regulated, immediate early genes in the innate response are thought to be primarily controlled at the level of transcriptional elongation. In uninfected cells, innate responsive genes are inactive, but found in open chromatin domains engaged by inactive/hypophosphorylated RNA Pol II [56, 57]. Upon infection, NF*κ*B repositions activated BRD4 complex. BRD4 kinase activity phosphorylates cyclin-dependent kinases and the carboxyterminal domain of RNA Pol II [58], transitioning it into an elongation-competent mode to express fully spliced inflammatory and antiviral genes.

Our study substantially extends this model by demonstrating that a subset of innate and stress-response genes undergo transcriptional elongation-coupled alternative splicing. These functional results are consistent with our earlier findings that the spliceosome binds to BRD4 in RSV infection, and is dependent on BD interaction with acetylate lysines. Future studies will be required to identify the spliceosome factors important for *IFRD1* and *ATP11A* isoform switching.

### 3.2. BRD4 inhibitors as a tool to study bromodomain biology

Bromodomain-and-extra-terminal-domain (BET) inhibitors have been used extensively over the last decade to probe bromodomain protein biology and disrupt related transcriptional activity [59, 60]. Small molecule BET inhibitors function by competitively occupying the bromodomains of this family of proteins, and significant effort has been dedicated to developing protein-specific inhibitors for targeted inhibition and degradation of individual BET proteins. In the context of airway inflammation, BET inhibition has been shown to block BRD4-mediated inflammatory activation and induced expression of cytokines and interferons [61]. Furthermore, BET inhibition blocks airway fibrosis and remodelling that proceeds through the epithelial-to-mesenchymal transition (EMT).

Classically, bromodomain inhibitors suffer from low specificity and high toxicity [60]. To address this issue and specifically probe BRD4, we use the highly specific and selective BRD4 inhibitor ZL0454, previously developed and used by our research group [29], using structure-based drug design to identify acetyl-lysine blocking chemistries. ZL0454 shows high selectivity for both bromodomains of BRD4, and displaces acetylated lysine side chains from the bromodomain (BD)-1 and −2 of BRD4 with an IC50 of approximately 50 nM as determined by time-resolved fluorescence resonance energy transfer (TR-FRET) assays. Furthermore, ZL0454 binds to BRD4 16-20 times stronger than the bromodomains of the closely related BRD2, BRD3, and BRDT. Compared to other notable BET inhibitors, ZL0454 also does not produce detectable toxicity in cell culture or in vivo. Finally, ZL0454 has been shown to displace interactors from the BRD4 complex, including key members of the spliceosome [12]. These traits make ZL0454 an ideal probe to study the role of BRD4 at the confluence of alternative splicing and innate inflammation.

### 3.3. BRD4 regulates alternative splicing of key innate inflammation coregulators

Recent works have demonstrated that BRD4 interacts with components of the spliceosome, and that these interactions can mediate alternative splicing in the context of T cell acute lymphoblastic leukemia [13]. Furthermore, BRD4:Spliceosome interactions have been shown to be dynamic in the context of airway viral infection, and sensitive to bromodomain inhibition [12]. In this work, we further investigate BRD4’s role in alternative splicing in the context of viral infection using a highly selective BRD4-specific inhibitor, ZL0454. In our data, treatment with ZL0454 resulted in detectable splicing events in 822 genes. Gene Ontology analysis identified only four significantly enriched pathways among these genes, suggesting that BRD4’s impact on alternative splicing in the airway epithelium is not biased for individual biological pathways, but rather widespread throughout the transcriptome. Nevertheless, observed splicing changes were significantly enriched for intron retention and extended Open Reading Frames, suggesting a coordinated mechanism of action. Among these alternatively spliced genes, we highlight three notable genes associated with innate inflammation, cytokine activation, and fibrosis: X-Box Binding Protein 1 (XBP1), ATPase Phospholipid Transporting 11A (ATP11A), and Interferon-related Developmental Regulator 1 (IFRD1).

XBP1 is a transcription factor essential to the activation of the immune response and the unfolded protein response [33, 36]. In the context of endoplasmic reticulum (ER) stress, the dual-function protein kinase/endoribonuclease IRE1*α* splices out a 26-nucleotide sequence from full-length XBP1 mRNA [34]. The resulting frame-shift enables translation of the active XBP1s protein, which translocates to the nucleus and serves as a potent transcriptional activator. In viral-induced innate inflammation, XBP1s directly binds the IL6 promoter and regulates expression of the key cytokine [36]. Furthermore, XBP1s has been implicated as a driver of fibrotic disease [62–64]. Our results further demonstrate that BRD4 regulates both the gene expression and alternative splicing of XBP1s, with IRE1*α* similarly effected, and provides a direct link between BRD4 and XBP1-mediated cytokine activation. However, the mechanism of this behavior, and the nature of its relationship with IRE1*α* remain to be fully elucidated. As BRD4 has been shown to interact with numerous transcription factors and chromatin remodelling factors [12], we propose that BRD4 may be serving as a scaffold for transcriptional co-regulators that directly control IRE1*α* and XBP1 gene expression. Future works that evaluate BRD4’s role in stabilizing such interactions at these gene bodies will be invaluable for understanding this mechanism of innate immune regulation.

ATP11A in contrast, is a membrane bound flippase - a class of enzymes responsible for ”flipping” phospholipids across surfaces of the cellular plasma membrane [39]. In particular, ATP11A is a member of the P4 ATPase Flippase complex, which has been directly linked to the recycling of Toll-like Receptor 4 (TLR4) back into the plasma membrane [40]. In human macrophages, knockdown of ATP11A induced hypersensitivity to lipopolysaccharide (LPS), due to an increase in membrane-associated TLR4. In our data set, we observed that BRD4i specifically reduces the expression of non-coding transcripts, while leaving protein-coding transcripts largely untouched, and in fact increased the abundance of ATP11A protein. This discrepancy likely signals the existence of additional layers of regulation, such as proteolytic degradation, but suggests that BRD4 may influence the activity of upstream components of the NF-*κ*B and innate immune signaling cascade. Additional studies to measure the cellular localization of TLR4 and other Toll-like receptors as a consequence of BRD4 inhibition will likely be illuminating.

Finally, we remark on IFRD1, which is especially interesting given its close relationship to NF-*κ*B p65/RelA. Studies in muscle tissue and keratinocytes have demonstrated that IFRD1 associates with histone deacetylases 3 and 4 (HDAC3/HDAC4) to attenuate the acetylation state of RelA [37, 65]. As RelA acetylation is a key marker of innate inflammation and is required for its interaction with BRD4 and the pTEFb complex [6, 27], this led to an expected decrease in the abundance of pro-inflammatory cytokines. IFRD1 overexpression in mice and human bronchiole epithelioid cells also reduced inflammation [66]. However, neutrophils from IFRD1-knockout mice demonstrate a reduced inflammatory phenotype, and our data suggests that IFRD1 in small airway epithelial cells actually blunts basal inflammation [67]. These findings suggest cell-type and/or stimulus-dependent roles for IFRD1, and additional studies which directly examine IFRD1’s interactions with HDACs may be valuable to precisely qualify IFRD1’s role in airway inflammation. In addition, there is currently no high-sensitivity method to examine sitespecific RelA acetylation, with the exception of the K310Ac mark [68]. Novel, mass spectrometry-based analytical methods may enable precise quantitation of IFRD1’s impact on RelA acetylation, stratified by airway cell-type and stimulus.

### 3.4. BRD4 regulates the abundance of interacting splicing factors in airway epithelial cells

In the process of measuring protein-level splicing consequences, we discovered that BRD4 regulates the protein-level abundance of several splicing factors. In particular, we highlight the core splicing factor, U2AF subunit 2 (U2AF2), which is known to complex with RNA polymerase II [69], coupling alternative splicing to transcription. U2AF2 has also been found to interact directly with BRD4 [13], and is dynamically recruited by RSV-infection [12], suggesting a close, 3-way association between innate-activated transcriptional machinery and the spliceosome. Given the close interactions between these proteins, this finding initially suggested a post-translational mechanism for abundance regulation. However, targeted q-RT-PCR assays indicate that this regulation occurs at the transcript-level. The mechanism for this discrepancy is unclear, and may stem from sequencing artifacts or violation of standard differential gene expression assumptions [49–51]. However, this result agrees with our group’s previous finding - that BRD4 transcriptionally regulates the abundance of its interacting co-activators - and emphasizes the multifaceted, key role that BRD4 takes in systems biology [28]. At present, there is no ideal approach to deconvolute this extreme complexity, but multi-omics methodologies hold great promise to address these challenges as technology and methodologies continue to evolve.

### 3.5. Challenges in Multi-omic Analysis

Short-read RNA-sequencing is a widely applied technique for profiling changes to gene-product abundance in biological samples [50, 70]. When transcripts are summarized to gene-level statistics, RNA-seq is highly accurate and can achieve nearly complete genome coverage. However, it is well-established that transcript-level abundance and relative abundance changes do not correlate perfectly to protein-level abundance [71–73]. This owes to the multiple post-transcriptional mechanisms of abundance control, including alternative mRNA splicing and post-translational degradation. In this study, we address this limitation by utilizing a splice-aware analysis pipeline to measure transcript-level abundances, and high-sensitivity, liquid-chromatography - Data-Independent Analysis Parallel Acquisition - Serial Fragmentation Tandem Mass Spectrometry (LC-diaPASEF-MS/MS) to measure protein-level abundances.

This integrated analysis offered combined coverage of ~7500 genes, however, it is worth noting that the quantifiable transcriptome of hSAECs includes nearly 20,000 genes. Many proteins, such as XBP1, are present at such low abundances that they cannot currently be quantified by our method. Significant improvements in proteomics technologies, particularly focused towards sensitivity and dynamic range, will be essential for comprehensive multi-omic quantitation of gene products. Furthermore, both short-read RNA-sequencing and shotgun proteomics use short, fragmented sequences as proxy for whole-transcript/protein abundance [70, 74]. This approach, while robust for overall quantitation, can not comprehensively identify and quantify all possible transcript variants and proteoforms [75]. Recent advances in long-read nano-pore sequencing [76, 77], and future advances in intact protein analysis [78–80] may allow for a truly comprehensive multi-omic analysis pipeline.

### 3.6. Conclusions

In summary, we utilize multi-omics approach integrating state-of-the-art proteomics and transcriptomic profiling, in concert with BRD4-specific small molecule bromodomain inhibitors, to identify BRD4-mediated alternative splicing in the context of the human airway epithelium. Further interrogating the results, we link BRD4-mediated alternative splicing to key regulators of innate activation, including X-box Binding Protein 1 (XBP1), ATPase Phospholipid Transporting 11A (ATP11A), and Interferon-related Developmental Regulator 1 (IFRD1). Finally, functional analysis of IFRD1 knockdown in this cell line revealed unexpected negative regulation of cytokine expression. These results directly clarify mechanisms of BRD4-mediated innate inflammation.

## Supporting information

Key Resources Table

## 4. Acknowledgements

The authors would like to acknowledge the UW-Madison Human Proteomics Program Mass Spectrometry Facility (initially funded by the Wisconsin partnership funds) for support in obtaining mass spectrometry data and NIH S10OD018475 for the acquisition of ultra-high resolution mass spectrometer for biomedical research. We would also like to thank Bruker for providing the timsTOF Pro mass spectrometer that was used in the study for the collection of all mass spectrometry data.

This work was partially supported by NIH grants AI062885 (ARB) and NCATS UL1TR002373 (ARB). D.S.R. would like to acknowledge support from the American Heart Association Predoctoral Fellowship Grant No. 832615/David S. Roberts/2021. J.Z. is also partly supported by the John D. Stobo, M.D. Distinguished Chair Endowment Fund. The funders had no role in the design of the study; in the collection, analyses, or interpretation of data; in the writing of the manuscript, or in the decision to publish the results.

## 5. Author Contributions

**Morgan Mann:** Conceptualization, Methodology, Software, Validation, Formal Analysis, Investigation, Writing - Original Draft, Visualization. **Yao Fu:** Validation, Investigation, Writing - Review & Editing. **Xiaofang Xu:** Resources, Writing - Review & Editing. **David S. Roberts:** Resources, Writing - Review & Editing, Supervision. **Yi Li:** Resources, Writing - Review & Editing. **Jia Zhou:** Resources, Writing - Review & Editing. **Ying Ge:** Resources, Supervision, Writing - Review & Editing, Project Administration, Funding Acquisition. **Allan R. Brasier:** Conceptualization, Resources, Writing - Review & Editing, Supervision, Project Administration, Funding Acquisition.

## 8. Methods

### 8.1. RESOURCE AVAILABILITY

#### 8.1.1. Lead Contact

Further information and requests for resources and reagents should be directed to and will be fulfilled by the lead contact, Allan R. Brasier.

#### 8.1.2. Materials Availability

This study did not generate new unique reagents.

#### 8.1.3. Data Availability Statement

- This paper analyzes existing, publicly available data. These accession numbers for the datasets are listed in the key resources table. Raw mass spectrometry data have been deposited to the ProteomeXchange Consortium via the PRIDE [81] partner repository, and are publicly available as of the date of publication. DOIs are listed in the key resources table.
- No original code was generated in this study.
- Any additional information required to reanalyze the data reported in this paper is available from the lead contact upon request.

### 8.2. EXPERIMENTAL MODEL & SUBJECT DETAILS

#### 8.2.1. Virus Preparation

The human RSV long strain was grown in Hep-2 cells and prepared by sucrose gradient centrifugation as previously described [6, 82–84]. The viral titer of purified RSV pools was varied from 8 to 9 log PFU/ml, determined by a methylcellulose plaque assay [84, 85]. Viral pools were aliquoted, quick-frozen on dry ice-ethanol, and stored at −70 °C until used.

#### 8.2.2. Cell Culture

Primary human small airway epithelial cells (hSAECs) were immortalized using human Telomerase/CDK4 as previously described [86, 87], and grown in SAGM small airway growth medium (Lonza, Walkersville, MD, USA). All cells were incubated at 37 °C, 5% CO_2_ until confluence.

A549 alveolar epithelial cells were purchased from the ATCC, and grown in F12K medium supplemented with 10% fetal bovine serum.

The BRD4-selective BD competitive inhibitor, ZL0454, was synthesized as synthesized as previously described [29, 61], and determined to be > 99% pure. ZL0454 was dissolved in dimethylsulfoxide (DMSO) and added to the relevant cell culture media at a final concentration of 10 *μ*M. The ZL0454 inhibitor or vehicle (DMSO) was added 18 hours before infection/stimulation, and to the media during infection/stimulation. hSAECs were infected with viral particles at a multiplicity of infection (MOI) of 1, or left uninfected (Mock). hSAECs were harvested at 24 hours post-infection.

Polyinosinic-polycytidilyic acid (polyIC) () was reconstituted in phosphate-buffered saline (PBS) at a concentration of 10 mg/ml. Reconstituted polyIC was added to cell culture medium at a final concentration of 50 *μ*g/ml, and consequently added to hSAECs for stimulation. PBS was used as a vehicle control. hSAECs were stimulated with polyIC for 15, 30, or 60 minutes before harvest.

#### 8.2.3. siRNA Knockdown

Non-targeting (Cat. # T-2001-01) and IFRD1-specific (Cat. # L-019615-00-0005) ON-TARGETplus SMARTPool siRNA was purchased from Horizon Discovery Biosciences (Cambridge UK). siRNA was transfected into hSAECs as per the manufacturer’s instructions, and cells were harvested for protein and RNA 72 hours later.

### 8.3. METHOD DETAILS

#### 8.3.1. Reagents & Chemicals

The 4-hexylphenylazosulfonate (Azo) used in these experiments was synthesized in-house as described previously [88, 89]. The BRD4 selective BD competitive inhibitor, ZL0454, was synthesized as previously described [29, 61] and determined to be >99% pure.

#### 8.3.2. Proteomics Sample Preparation

Human small airway epithelial cells (hSAECs) were lysed in a buffer containing 0.2% Azo, 25mM Ammonium Bicarbonate, 10 mM L-Methionine, 1 mM Dithiothreitol (DTT), 1x HALT Protease and Phosphatase Inhibitor (Thermofisher, Waltham, MA, USA, Cat. # 78440), before transfer to microcentrifuge tubes and boiling at 95 °C for 5 minutes. Samples were diluted to 0.1% Azo, and standardized to 0.5 mg/ml by Bradford Assay (Bio-Rad, Hercules, CA, USA, Cat. # 5000006), prior to chemical reduction using 30 mM DTT for 60 minutes at 37 °C. Freshly prepared iodoacetamide solution (200 mM) was added to a final concentration of 20 mM, and the samples were incubated in the dark for 30 min. Protein was digested with Trypsin Gold (Promega, Madison, WI, USA) at a 1:50 enzyme:protein ratio overnight at 37 °C with agitation at 1000 rpm.

Digestion was quenched by addition of formic acid to a final concentration of 1%, and irradiated at 305 nM for 5 minutes to photocleave the Azo surfactant. Samples were centrifuged at 15,000xg, and supernatents were desalted using Pierce C18 tips (Thermo Scientific, Waltham, MA, USA). Peptide pellets were resuspended in 0.2% Formic Acid immediately prior to LC-MS analysis.

#### 8.3.3. Online Data Independent Analysis - Parallel Accumulation Serial Fragmentation (diaPASEF) Label-Free Quantitative Proteomics

Desalted peptides (200 ng) were loaded and separated on an IonOptiks Aurora UHPLC column with CSI fitting (Melbourne, Australia) at a flow rate of 0.4 *μ*L/min and a linear gradient increasing from 0% to 17% mobile phase B (0.1% formic acid in acetonitrile) (mobile phase A: 0.1% formic acid in water) over 60 min; 17% to 25% from 60 to 90 min; 25% to 37% B from 90 to 100 min; 37% to 85% B from 100 min to 110 min; and a 10 min hold at 85% B before washing and returning to low organic conditions. The column directly integrated a nanoESI source for delivery of the samples to the mass spectrometer. MS spectra were captured with a Bruker timsTOF Pro quadrupole-time of flight (Q-TOF) mass spectrometer (Bruker Daltonics, Billerica, MA, USA) operating in diaPASEF mode, using 32 windows ranging from m/z 400 to 1200 and 1/K0 0.6 to 1.42. Precise windows settings are available in the Table S4.

#### 8.3.4. RelA Immunoprecipitation

A549 alveolar epithelial cells were lysed in Low Ionic Strength Immunoprecipitation Buffer (50 mM NaCl, 10 mM HEPES, 1% Triton-X100, 10% Glycerol) with 1 mM DTT and 1% Protease Inhibitor Cocktail (Millipore-Sigma, Burlington, MA, USA, Cat. # P8340), and extracts were sonicated 3x for 10 seconds each time (BRANSON Sonifier 150, setting 4) prior to centrifugation at 10000xg, 4 °C for 10 minutes. Supernatants were collected, and total protein abundance was normalized to 2 *μg* using the Detergent-compatible Protein Assay (Biorad, Hercules, CA, USA, Cat. # 5000111). 2 *μ*g *α*-RelA antibody (Cell Signaling, Danvers, MA, USA, Cat. # 8242) was added to each extract, and the samples were incubated overnight at 4 °C with rotation.

The next day, 30 *μ*l Protein G-conjugated magnetic beads (Dynabeads, Invitrogen, Waltham, MA, USA, Cat. # 10003D) were added to each sample and allowed to bind for 4 hours at 4 °C. After binding, supernatents were removed, and the beads were washed 3x in Low Ionic Strength Immunoprecipitation Buffer, prior to elution by boiling in 2x Laemli loading buffer (BioRad, Hercules, Ca, USA, Cat. # 1610737EDU) at 95 °C for 5 minutes. Eluate was examined by western blot, as detailed in the next section.

#### 8.3.5. Western Blotting

For total protein quantitation, cells were lysed and protein extracted using radio-immunoprecipitation buffer (50 mM Tris-HCL (pH 7.6), 150 mM NaCl 1% NP-40, 0.5% Sodium Deoxycholate, 0.1% SDS) with 1 mM Dithithreitol and 1x Protease Inhibitor Cocktail (MilliporeSigma, Burlington, MA, USA, Cat. # P8340) added. Protein was normalized using the Detergent-compatible Protein Assay (Biorad, Hercules, CA, USA, Cat. # 5000111), and loaded onto a 4–20% Criterion TGX Precast Protein Gel (Biorad, Hercules, CA, USA) for separation. Immunoprecipitation samples were prepared as detailed in the previous section.

Proteins were transferred to a nitrocellulose membrane using a Trans-Blot Turbo Transfer System (Biorad, Hercules, CA, USA) with a constant voltage of 25 V over 30 min. The membrane was blocked for 1 h using 5% milk powder in Tris-buffered Saline with 0.1% Tween-20 (TBST) and incubated overnight at 4 °C with primary antibody (see Table **??**) diluted in 5% milk powder in TBST. Membranes were washed thoroughly and incubated with a horseradish peroxidase (HRP)-conjugated secondary antibody diluted 1:10000 in 5% milk powder in TBST for 1 hour at room temperature. The VeriBlot IP Detection Reagent (Abcam, Cambridge, UK, Cat. # ab131366) was used as a secondary antibody for immunoprecipitated samples. Imaging was performed via chemiluminescent detection using Supersignal West Femto Maximum Sensitivity Substrate (ThermoFisher, Waltham, MA, USA, Cat. # 34095) and an Azure c500 gel imaging system (Azure Biosystems, Dublin, CA, USA). When necessary, bands were quantified by densitometry, using FIJI version 1.53c [90]. Membranes were stripped between incubation with different antibodies using Restore Western Blot Stripping Buffer (ThermoFisher, Waltham, MA, USA, Cat. # 21059).

#### 8.3.6. RNA Isolation & Reverse-Transcriptase PCR

Cellular RNA from from cultured cells was isolated using an RNeasy kit with on-column DNase digestion (Qiagen, Germantown, MD, USA, Cat. # 74004). RNA was reverse transcribed into cDNA using the First Strand cDNA Synthesis kit (Thermo Fisher Scientific). Quantitative PCR (qPCR) reactions were conducted in duplicate using SYBR Green Master mix (Bio-Rad) and 500 ng of cDNA in 20 ul reactions. Gene-specific primers were used at a concentration of 500 nM and are listed below. The wPCR reaction was carried out on a AriaMx Real-Time PCR System (Agilent, Santa Clara, CA, USA) with 40 cycles of 15 s at 95 °C and 30 s at 60 °C. Cyclophilin A (PP1A) was used as a housekeeping gene for overall gene expression. XBP1-Total was used to normalize the abundance of spliced XBP1 (XBP1-Short).

IFRD1 splicing variants were amplified using Q5 High Fidelity Polymerase (New England Biolabs, Ipswich, MA, USA, Cat. # M0491S) on an AriaMX Real-Time PCR System (Agilent, Santa Clara, CA, USA) with 35 cycles of 10 s at 98 °C, 30 s at 67 °C, and 30 s at 72 °C. Due to the significant heterogeneity of IFRD1 transcript variants, primers for total IFRD1 (IFRD1-Total) were chosen amplify a common region, bridging exons 7 and 8, found in all proteincoding transcripts. The expected amplification products (IFRD1-204: 707, IFRD1-Total: 201 bp) were imaged via UV detection on an Azure c500 gel imaging system (Azure Biosystems, Dublin, CA, USA), and quantified by Gel Densitometry.

#### 8.3.7. Two-step Crosslinking Chromatin Immunoprecipitation (XChIP) - PCR

Cells were cross-linked in-vivo using disucinimidylglutarate (DSG) and methanol-free formaldehyde as previously described [91]. After lysis (1% SDS, 50 mM Tris-HCl, pH 8.0), extracts were sonicated 3x for 10 seconds each time (BRANSON Sonifier 150, setting 4), and centrifuged for 5 min at 10,000 g, 4 °C. Supernatents were diluted 10-fold in Low Ionic Strength Immunoprecipitation Buffer (50 mM NaCl, 10 mM HEPES, 1% Triton-X100, 10% Glycerol) with 1 mM DTT and 1% Protease Inhibitor Cocktail (MilliporeSigma, Burlington, MA, USA, Cat. # P8340). Constant volumes of sample were incubated overnight with 2 *μ*g *α*-BRD4 antibody (Cell Signaling, Danvers, MA, USA, Cat. # 13440) or a nonspecific isotype control (LSBio, Seattle, WA, USA, Cat. # LS-C149375).

The next day, 20 *μ*l Protein G-conjugated magnetic beads (Dynabeads, Invitrogen, Waltham, MA, USA) were added to each sample and incubated for 1 hour at 4 °C. 50 *μ*l aliquots were taken from each sample to serve as Input controls. All samples were washed twice in Low Ionic Strength IP buffer, once in High Ionic Strength Wash Buffer (500 mM NaCl, 0.1% SDS, 1% Triton-X100, 2 mM EDTA, 20 mM Tris-HCl, pH 8.0), and twice in Tris-EDTA Buffer (10 mM Tris-HCl, 1 mM EDTA, pH 8.0) prior to elution in 300 *μ*l 0.1 M Sodium Bicarbonate, 1% SDS at 65 °C. Samples were decrosslinked via the addition of 20 *μ*g Proteinase K, and incubated overnight at 65 °C.

Decrosslinked DNA was purified by Phenol-Chloroform Extraction and Ethanol Precipitation. In brief, 100 *μ*l of 5:1 Phenol:Chloroform was added to each sample prior to vortexing, and centrifugation at 14000xg, 24 °C for 10 minutes. The supernatent was transferred to a new tube containing 100 *μ*l of chloroform, and vortexed and centrifuged again as above. Finally, the supernatents (~200 *μ*l) were transferred to a new tube containing 100 *μ*l 7.5 M Ammonium Acetate and 60 *μ*g of glycogen. 1 ml of 100% ethanol was added to each tube, and the samples were vortexed prior to overnight precipitation at −80 °C. Purified DNA was reconstituted with DNase-free water, and quantified by qPCR. Input control samples were used for normalization.

### 8.4. Quantification & Statistical Analysis

#### 8.4.1. Short-Read RNA Sequencing & Splicing Analysis

RNA-Sequencing (RNA-Seq) data was obtained *Xu et. al.* [28]. The data was analyzed using R version 4.1.0 and the ”IsoformSwitchAnalyzeR” R Package [92], utilizing Gencode version 33 transcripts [93], and integrating the ”CPAT” [94], ”PFAM” [95], ”SignalP” [96], and ”IUPRED2A” [97] tools to characterize transcripts based on coding status and switch consequence. Isoform switches were identified using an isoform switch q-value threshold of 0.05 and an isoform fraction difference of 0.1. Significant results were submitted to PANTHER gene list statistical over-representation test, using ”PANTHER Biological Process Complete” as the annotation terms, and the Homo Sapiens genome as the reference list [31, 32]. 2-Dimensional Annotation analysis was carried out in Perseus [98] as described in *Cox & Mann* [48], using Kyoto Encycolopedia of Genes and Genomes (KEGG) pathways [99] as annotations. Gene ontology and annotation terms were filtered to a 1% false discovery rate (FDR). Figures were generated using the ”ggplot2” [100] and ”ggpubr” [101] R packages for R version 4.1.0 [102]. Boxplots represents the median and interquartile range of n=4 biological replicates.

#### 8.4.2. Protein-level Differential Abundance Analysis

Raw LC-MS data was quantified using DIA - Neural Network (DIANN) version 1.8,[103] using the following parameters: 1% FDR, Library-free search enabled, Minimum fragment m/z: 200, Maximum fragment m/z: 1800, Minimum precursor m/z: 400, Maximum precursor m/z: 1200, Minimum precursor charge: 2, Maximum precursor charge: 4, Minimum peptide length: 7, maximum peptide length: 30, Enzyme: Trypsin, N-terminal methionine cleavage enabled, cysteine carbamidomethylation enabled, Maximum missed cleavages: 1, MS1/MS2 mass accuracy: 10 ppm, Quantification strategy: Robust LC (High Precision), Neural network classifier: Double-pass mode. All other settings were left at default values. Data was searched against a fasta-formatted text file containing 20,404 reviewed human protein sequences (Taxon ID: 9606) and 6 sequences from the RSV Long-strain (Taxon ID: 11260) [104].

Protein-level quantification data was filtered using the ”DAPAR” package [105] for R version 4.1.0 to include all proteins quantified in 2 of 3 runs in at least one sample group. Values were then median normalized and missing values were imputed via ssla for partially observed values within a condition, or set to the 2.5% quantile of observed intensities for observations that were missing entirely within a condition. The ”DEP” R package [106] was used to perform a Limma test between all specified contrasts, and the ”IHW” R package [107] was used to adjust all p-values, using the number of quantified peptides per protein as a covariate. A PAdj threshold of 0.05 and a Log_2_ Fold Change threshold of 0.6 were set to identify significant changes to protein abundance. Tandem MS spectra were visualized and plotted using Skyline [108] and the “ggplot2” [100] package for R version 4.1.0 [102]. Boxplots were generated using the “ggpubr” [101] package for R version 4.1.0 [102].

#### 8.4.3. Gel Densitometry

Gel bands were quantified using FIJI version 1.53c [90], and statistical analysis was conducted by One- or Two-Factor ANOVA, with post-hoc Tukey’s HSD. Boxplots were generated using the “ggpubr” [101] package for R version 4.1.0 [102]. Boxplots represent median and interquartile range of n=3 biological replicates. Barplots represent mean ± standard error (SE) of n=3 biological replicates.

#### 8.4.4. qPCR Quantification & Analysis

Data was analyzed using the △△Ct method. Statistical analysis was conducted by Two-Factor ANOVA and post-hoc Tukey’s HSD. Boxplots were generated using the “ggpubr” [101] package for R version 4.1.0 [102], and represent median and interquartile range of n=3 biological replicates. Scatter plots represent mean ± standard error (SE) of n=3 biological replicates.

### 8.5. KEY RESOURCES TABLE

Key Resources Table is provided as a microsoft word file in the requested format.

## 9. Supplementary Information Titles & Legends

**Figure S1.**
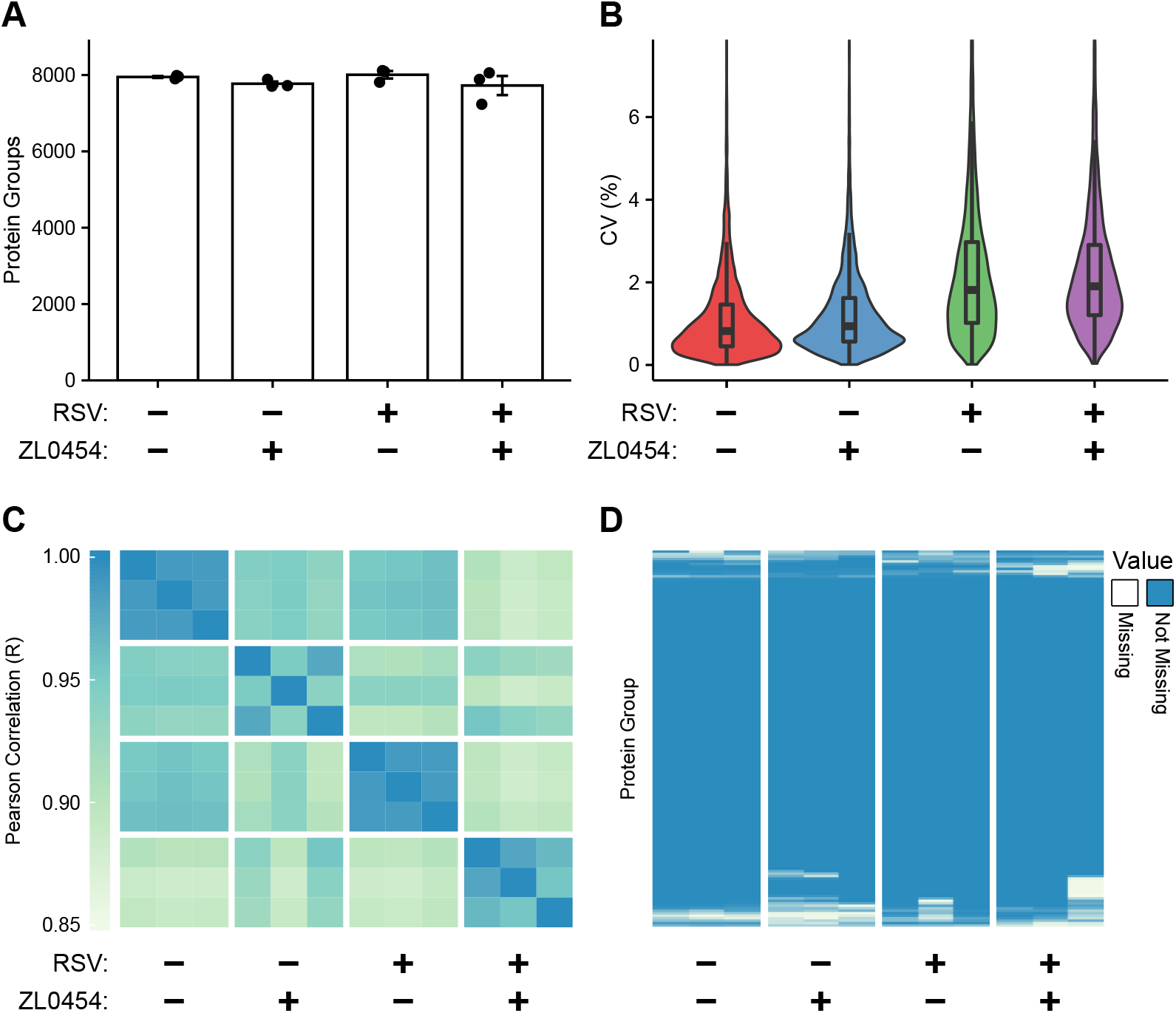
Quality control metrics for diaPASEF LCMS analysis. (A) Quantifiable proteins per experimental group. (B) Coefficient of Variance (%CV) violin plots indicating intra-group median variance to protein abundance measurements. (C) Heatmap indicating sample-to-sample Pearson Correlation scores. (D) Data completeness matrix of quantifiable proteins. Data is prior to filtering.

**Figure S2.**
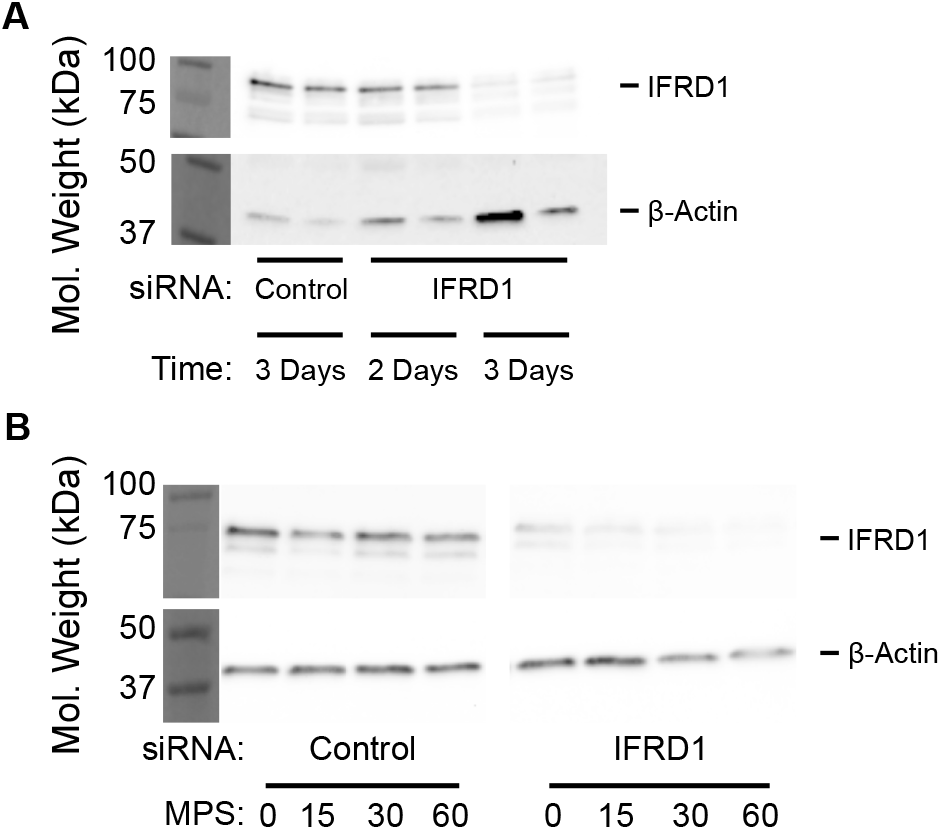
Optimization of IFRD1 knockdown by siRNA in hSAECs. (A) Western blot indicating peak protein knockdown occurs at 72+ hours. (B). Western blot demonstrating that IFRD1 protein abundance is stable throughout inflammatory activation by polyIC after treatment with either non-targetting or IFRD1-specific siRNA.

**Table S1.** Isoform Switch Results Table from IsoformSwitchAnalyzer. Table included in attached .xlsx file.

**Table S2.** Protein differential abundance results. Table included in attached .xlsx file.

**Table S3.** Raw DIA-NN output indicating protein quantification values by replicate. Table included in attached .xlsx file.

**Table S4.** diaPASEF Window Parameters. Table included in attached .xlsx file.

**Table S5.** Oligonucleotide Sequences. Table included in attached .xlsx file.

