## Supplementary material for "Bromodomain-containing Protein 4 Regulates Innate Inflammation in Airway Epithelial Cells via Modulation of Alternative Splicing": Key Resources Table

The table highlights the reagents, genetically modified organisms and strains, cell lines, software, instrumentation, and source data **essential** to reproduce results presented in the manuscript. Depending on the nature of the study, this may include standard laboratory materials (i.e., food chow for metabolism studies, support material for catalysis studies), but the table is **not** meant to be a comprehensive list of all materials and resources used (e.g., essential chemicals such as standard solvents, SDS, sucrose, or standard culture media do not need to be listed in the table). **Items in the table must also be reported in the method details section within the context of their use.** To maximize readability, the number of **oligonucleotides and RNA sequences** that may be listed in the table is restricted to no more than 10 each. If there are more than 10 oligonucleotides or RNA sequences to report, please provide this information as a supplementary document and reference the file (e.g., See Table S1 for XX) in the key resources table.

***Please note that ALL references cited in the key resources table must be included in the main references list.*** Please report the information as follows:

- **REAGENT or RESOURCE:** Provide the full descriptive name of the item so that it can be identified and linked with its description in the manuscript (e.g., provide version number for software, host source for antibody, strain name). In the experimental models section (applicable only to experimental life science studies), please include all models used in the paper and describe each line/strain as: model organism: name used for strain/line in paper: genotype. (i.e., Mouse: OXTR^fl/fl^: B6.129(SJL)-Oxtr^tm1.1Wsy/J^). In the biological samples section (applicable only to experimental life science studies), please list all samples obtained from commercial sources or biological repositories. Please note that software mentioned in the methods details or data and code availability section needs to also be included in the table. See the sample tables at the end of this document for examples of how to report reagents.
- **SOURCE:** Report the company, manufacturer, or individual that provided the item or where the item can be obtained (e.g., stock center or repository). For materials distributed by Addgene, please cite the article describing the plasmid and include “Addgene” as part of the identifier. If an item is from another lab, please include the name of the principal investigator and a citation if it has been previously published. If the material is being reported for the first time in the current paper, please indicate as “this paper.” For software, please provide the company name if it is commercially available or cite the paper in which it has been initially described.
- **IDENTIFIER:** Include catalog numbers (entered in the column as “Cat#” followed by the number, e.g., Cat#3879S). Where available, please include unique entities such as [RRIDs](https://www.force11.org/group/resource-identification-initiative), Model Organism Database numbers, accession numbers, and PDB, CAS, or CCDC IDs. For antibodies, if applicable and available, please also include the lot number or clone identity. For software or data resources, please include the URL where the resource can be downloaded. Please ensure accuracy of the identifiers, as they are essential for generation of hyperlinks to external sources when available. Please see the Elsevier [list of data repositories](https://www.elsevier.com/authors/author-resources/research-data/data-base-linking) with automated bidirectional linking for details. When listing more than one identifier for the same item, use semicolons to separate them (e.g., Cat#3879S; RRID: AB_2255011). If an identifier is not available, please enter “N/A” in the column.
  - ***A NOTE ABOUT RRIDs:*** We highly recommend using RRIDs as the identifier (in particular for antibodies and organisms but also for software tools and databases). For more details on how to obtain or generate an RRID for existing or newly generated resources, please [visit the RII](https://www.force11.org/group/resource-identification-initiative) or [search for RRIDs](https://scicrunch.org/resources).

Please use the empty table that follows to organize the information in the sections defined by the subheading, skipping sections not relevant to your study. Please do not add subheadings. To add a row, place the cursor at the end of the row above where you would like to add the row, just outside the right border of the table. Then press the ENTER key to add the row. Please delete empty rows. Each entry must be on a separate row; do not list multiple items in a single table cell. Please see the sample tables at the end of this document for relevant examples in the life and physical sciences of how reagents and instrumentation should be cited.

***TABLE FOR AUTHOR TO COMPLETE***

*Please upload the completed table as a separate document.* ***Please do not add subheadings to the key resources table.*** *If you wish to make an entry that does not fall into one of the subheadings below, please contact your handling editor.* ***Any subheadings not relevant to your study can be skipped.*** *(****NOTE:*** *References within the KRT should be in numbered style rather than Harvard.)*

**Key resources table**

| REAGENT or RESOURCE | SOURCE | IDENTIFIER |
| --- | --- | --- |
| Antibodies | | |
| Rabbit polyclonal anti-IFRD1 | Abcam | Cat. # ab229720 |
| Rabbit monoclonal anti -NF$\kappa$B p65 XP® | Cell Signaling Technology | Cat. # 8242 |
| Rabbit monoclonal anti- NF$\kappa$B p65 (Lys310) | Cell Signaling Technology | Cat. # 3045 |
| Rabbit monoclonal anti- Acetylated-lysine (Ac-K^2^-100) Multi-Mab^TM^ | Cell Signaling Technology | Cat. # 9814 |
| Mouse monoclonal anti-Beta Actin | Abcam | Cat. # ab8226 |
| Rabbit polyclonal IgG Isotype Control Antibody | LSBio | Cat. # LS-C149375 |
| Goat anti-Rabbit IgG (H+L) Secondary Antibody, HRP | ThermoFisher Scientific | Cat. # 31460 |
| Goat anti-Mouse IgG (H+L) Secondary Antibody, HRP | ThermoFisher Scientific | Cat. # 62-6520 |
| Veriblot for IP Detection Reagent (HRP) | Abcam | Cat. # ab131366 |
| Bacterial and virus strains | | |
| Human Respiratory Syncytial Virus (RSV, long strain) | ATCC | Cat. # VR-26 |
| Biological samples |  |  |
| Chemicals, peptides, and recombinant proteins | | |
| 4-hexylphenylazosulfonate | Brown, et. al. *Nat Methods.* **2019** | NA |
| Acetonitrile | MilliporeSigma | Cat. # 1000291000 |
| Ammonium Acetate | MilliporeSigma | Cat. # A1542-250G |
| Ammonium Bicarbonate | ThermoFisher Scientific | Cat. # 014249.A1 |
| Disucinimidyl glutarate (DSG) | ThermoFisher Scientific | Cat. # 20593 |
| Dithiothreitol (DTT) | MilliporeSigma | Cat. # 646563 |
| DMSO | MilliporeSigma | Cat. # D8418-50ML |
| DNase-free water | ThermoFisher Scientific | Cat. # 46-2224 |
| Ethanol (100%) | MilliporeSigma | Cat. # E7023-500ML |
| Formic Acid | MilliporeSigma | Cat. # 5330020050 |
| Glycogen | MilliporeSigma | Cat. # 10901393001 |
| Glycerol | MilliporeSigma | Cat. # 911038-500ML |
| HEPES | TEKNova | Cat. # H1090 |
| Hydrochloric Acid | LabChem | Cat. # LC153603 |
| Iodoacetamide | MilliporeSigma | Cat. # I6125-G |
| L-Methionine | MilliporeSigma | Cat. # 64319-100G-F |
| L-Histidine | MilliporeSigma | Cat. # H8000-100G |
| NP-40 Alternative | MilliporeSigma | Cat. # 492016-100ML |
| 5:1 Phenol:Chloroform | MilliporeSigma | Cat. # P1944 |
| Phosphate-buffered Saline (PBS) | MilliporeSigma | Cat. # P3813-10PAK |
| Sodium Deoxycholate | MilliporeSigma | Cat. # D6750-100G |
| Sodium Dodecylsulfate (10%) | Corning | Cat. # 46-040-CI |
| Sodium Bicarbonate | MilliporeSigma | Cat. # S5761-500G |
| Sodium Chloride (NaCl) | Dot Scientific Inc. | Cat. # DSS23020-500 |
| Tween-20 | MilliporeSigma | Cat. # P1379-250ML |
| Triton-X100 | MilliporeSigma | Cat. # X100-100ML |
| Tris-EDTA Buffer | ThermoFisher Scientific | Cat. # AM9858 |
| Tris Base | MilliporeSigma | Cat. # 252859-500G |
| ZL0454 | Tian, et. al. *Am. J. Respir Cell Mol Biol.* **2019** | NA |
| Blotting Grade Blocker Non Fat Dry Milk Powder | BioRad | Cat. # 1706404XTU |
| Dynabeads Protein G | ThermoFisher Scientific | Cat. # 10003D |
| Laemmli Sample Buffer | BioRad | Cat. # 1610737EDU |
| HALT Protease and Phosphatase Inhibitor | ThermoFisher Scientific | Cat. # 78440 |
| Protease Inhibitor Cocktail | MilliporeSigma | Cat. # 8340 |
| Restore Western Blot Stripping Buffer | ThermoFisher Scientific | Cat. # 21059 |
| Supersignal West Femto Maximum Sensitivity Substrate | ThermoFisher Scientific | Cat. # 34095 |
| PolyIC | MilliporeSigma | Cat. # P0913-10MG |
| Q5 High Fidelity Polymerase | New England Biolabs | Cat. # M0491S |
| Proteinase K | MilliporeSigma | Cat. # 70663-4 |
| Trypsin Gold | Promega | Cat. # V5280 |
| Critical commercial assays | | |
| Detergent-compatible Protein Assay | BioRad | Cat. # 5000111 |
| Bradford Protein Assay | BioRad | Cat. # 5000006 |
| RNeasy RNA Purification Kit | Qiagen | Cat. # 74004 |
| Superscript^TM^ III First Strand cDNA Synthesis Kit | ThermoFisher Scientific | Cat. # 18080051 |
| iTaq Universal SYBR Green Supermix | BioRad | Cat. # 1725121 |
| Deposited data | | |
| RNA-seq Data | GEO Database | Accession #: GSE179353 |
| Proteomics Data | ProteomeXchange | Project Accession: PXD039212 |
| Experimental models: Cell lines | | |
| Human Small Airway Epithelial Cells (hSAECs) | Ramirez, et. al. *Cancer Res.* **2004** | NA |
| Experimental models: Organisms/strains | | |
| Oligonucleotides | | |
| See Supplemental Table 3 |  |  |
| Recombinant DNA | | |
| Software and algorithms | | |
| R version 4.1.0 | R Core Team. R Foundation for Statistical Computing. **2022** | https://cran.r-project.org/bin/windows/base/old/ |
| IsoformSwitchAnalyzeR | Seerup & Sandelin. *Bioinformatics.* **2019** | https://bioconductor.org/packages/release/bioc/html/IsoformSwitchAnalyzeR.html |
| Gencode v33 Transcriptome | Frankish, et. al. *Nucleic Acids Res*. **2020** | https://www.gencodegenes.org/human/release_33.html |
| CPAT | Wang, et. al. *Nucleic Acids Res*. **2013** | https://bio.tools/cpat#:~:text=CPAT%20(Coding%2DPotential%20Assessment%20Tool,calculated%20from%20the%20RNA%20sequences. |
| PFAM | Mistry, et. al. *Nucleic Acids Res.* **2020** | http://pfam.xfam.org/ |
| SignalP | Armenteros, et. al. *Nat. Biotechnol*. **2019** | https://bio.tools/signalp |
| IUPRED2A | Erdos, et. al. *Curr Protoc Bioinformatics.* **2020** | https://iupred2a.elte.hu/ |
| PANTHER Gene Ontology | Thomas, et. al. *Genome Res.* **2003** | http://pantherdb.org/ |
| ggplot2 | Wichham, H. CRAN. **2016** | https://cran.r-project.org/web/packages/ggplot2/index.html |
| ggpubr | Kassambara, A. CRAN. **2020** | NA |
| DIANN | Demichev, et. al. *Nat. Methods.* **2020** | NA |
| DAPAR | Wieczorek, et. al. *Bioinformatics.* **2017** | NA |
| DEP | Zhang, et. al. *Nat. Protocols.* **2018** | NA |
| IHW | Ignatiadis, et. al. *Nat. Methods.* **2016** | NA |
| Skyline | Pino, et. al. *Mass Spectrom. Rev.* **2020** | NA |
| FIJI | Schindelin, et. al. *Nat. Methods.* **2012** | NA |
| Other | | |

***LIFE SCIENCE TABLE WITH EXAMPLES FOR AUTHOR REFERENCE***

| **REAGENT or RESOURCE** | **SOURCE** | **IDENTIFIER** |
| --- | --- | --- |
| Antibodies | | |
| Rabbit monoclonal anti-Snail | Cell Signaling Technology | Cat#3879S; RRID: AB_2255011 |
| Mouse monoclonal anti-Tubulin (clone DM1A) | Sigma-Aldrich | Cat#T9026; RRID: AB_477593 |
| Rabbit polyclonal anti-BMAL1 | This paper | N/A |
| Bacterial and virus strains | | |
| pAAV-hSyn-DIO-hM3D(Gq)-mCherry | Krashes et al.^1^ | Addgene AAV5; 44361-AAV5 |
| AAV5-EF1a-DIO-hChR2(H134R)-EYFP | Hope Center Viral Vectors Core | N/A |
| Cowpox virus Brighton Red | BEI Resources | NR-88 |
| Zika-SMGC-1, GENBANK: KX266255 | Isolated from patient (Wang et al.^2^) | N/A |
| *Staphylococcus aureus* | ATCC | ATCC 29213 |
| *Streptococcus pyogenes*: M1 serotype strain: strain SF370; M1 GAS | ATCC | ATCC 700294 |
| Biological samples | | |
| Healthy adult BA9 brain tissue | University of Maryland Brain & Tissue Bank; http://medschool.umaryland.edu/btbank/ | Cat#UMB1455 |
| Human hippocampal brain blocks | New York Brain Bank | http://nybb.hs.columbia.edu/ |
| Patient-derived xenografts (PDX) | Children's Oncology Group Cell Culture and Xenograft Repository | http://cogcell.org/ |
| Chemicals, peptides, and recombinant proteins | | |
| MK-2206 AKT inhibitor | Selleck Chemicals | S1078; CAS: 1032350-13-2 |
| SB-505124 | Sigma-Aldrich | S4696; CAS: 694433-59-5 (free base) |
| Picrotoxin | Sigma-Aldrich | P1675; CAS: 124-87-8 |
| Human TGF-β | R&D | 240-B; GenPept: P01137 |
| Activated S6K1 | Millipore | Cat#14-486 |
| GST-BMAL1 | Novus | Cat#H00000406-P01 |
| Critical commercial assays | | |
| EasyTag EXPRESS 35S Protein Labeling Kit | PerkinElmer | NEG772014MC |
| CaspaseGlo 3/7 | Promega | G8090 |
| TruSeq ChIP Sample Prep Kit | Illumina | IP-202-1012 |
| Deposited data | | |
| Raw and analyzed data | This paper | GEO: GSE63473 |
| B-RAF RBD (apo) structure | This paper | PDB: 5J17 |
| Human reference genome NCBI build 37, GRCh37 | Genome Reference Consortium | http://www.ncbi.nlm.nih.gov/projects/genome/assembly/grc/human/ |
| Nanog STILT inference | This paper; Mendeley Data | http://dx.doi.org/10.17632/wx6s4mj7s8.2 |
| Affinity-based mass spectrometry performed with 57 genes | This paper; Mendeley Data | Table S8; http://dx.doi.org/10.17632/5hvpvspw82.1 |
| Experimental models: Cell lines | | |
| Hamster: CHO cells | ATCC | CRL-11268 |
| *D. melanogaster*: Cell line S2: S2-DRSC | Laboratory of Norbert Perrimon | FlyBase: FBtc0000181 |
| Human: Passage 40 H9 ES cells | MSKCC stem cell core facility | N/A |
| Human: HUES 8 hESC line (NIH approval number NIHhESC-09-0021) | HSCI iPS Core | hES Cell Line: HUES-8 |
| Experimental models: Organisms/strains | | |
| *C. elegans*: Strain BC4011: [srl-1](http://www.wormbase.org/species/c_elegans/gene/WBGene00005636)([s2500](http://www.wormbase.org/search/variation/s2500)) II; [dpy-18](http://www.wormbase.org/species/c_elegans/gene/WBGene00001077)([e364](http://www.wormbase.org/search/variation/e364)) III; [unc-46](http://www.wormbase.org/species/c_elegans/gene/WBGene00006782)([e177](http://www.wormbase.org/search/variation/e177))[rol-3](http://www.wormbase.org/species/c_elegans/gene/WBGene00004395)([s1040](http://www.wormbase.org/search/variation/s1040)) V. | Caenorhabditis Genetics Center | WB Strain: BC4011; WormBase: WBVar00241916 |
| *D. melanogaster*: RNAi of Sxl: y[1] sc[*] v[1]; P{TRiP.HMS00609}attP2 | Bloomington Drosophila Stock Center | BDSC:34393; FlyBase: FBtp0064874 |
| *S. cerevisiae*: Strain background: W303 | ATCC | ATTC: 208353 |
| Mouse: R6/2: B6CBA-Tg(HDexon1)62Gpb/3J | The Jackson Laboratory | JAX: 006494 |
| Mouse: OXTRfl/fl: B6.129(SJL)-Oxtr^tm1.1Wsy^/J | The Jackson Laboratory | RRID: IMSR_JAX:008471 |
| Zebrafish: Tg(Shha:GFP)t10: t10Tg | Neumann and Nuesslein-Volhard^3^ | ZFIN: ZDB-GENO-060207-1 |
| *Arabidopsis*: 35S::PIF4-YFP, BZR1-CFP | Wang et al.^4^ | N/A |
| *Arabidopsis*: JYB1021.2: pS24(AT5G58010)::cS24:GFP(-G):NOS #1 | NASC | NASC ID: N70450 |
| Oligonucleotides | | |
| siRNA targeting sequence: PIP5K I alpha #1: ACACAGUACUCAGUUGAUA | This paper | N/A |
| Primers for XX, see Table SX | This paper | N/A |
| Primer: GFP/YFP/CFP Forward: GCACGACTTCTTCAAGTCCGCCATGCC | This paper | N/A |
| Morpholino: MO-pax2a GGTCTGCTTTGCAGTGAATATCCAT | Gene Tools | ZFIN: ZDB-MRPHLNO-061106-5 |
| ACTB (hs01060665_g1) | Life Technologies | Cat#4331182 |
| RNA sequence: hnRNPA1_ligand: UAGGGACUUAGGGUUCUCUCUAGGGACUUAGGGUUCUCUCUAGGGA | This paper | N/A |
| Recombinant DNA | | |
| pLVX-Tight-Puro (TetOn) | Clonetech | Cat#632162 |
| Plasmid: GFP-Nito | This paper | N/A |
| cDNA GH111110 | Drosophila Genomics Resource Center | DGRC:5666; FlyBase:FBcl0130415 |
| AAV2/1-hsyn-GCaMP6- WPRE | Chen et al.^5^ | N/A |
| Mouse raptor: pLKO mouse shRNA 1 raptor | Thoreen et al.^6^ | Addgene Plasmid #21339 |
| Software and algorithms | | |
| ImageJ | Schneider et al.^7^ | https://imagej.nih.gov/ij/ |
| Bowtie2 | Langmead and Salzberg^8^ | http://bowtie-bio.sourceforge.net/bowtie2/index.shtml |
| Samtools | Li et al.^9^ | http://samtools.sourceforge.net/ |
| Weighted Maximal Information Component Analysis v0.9 | Rau et al.^10^ | https://github.com/ChristophRau/wMICA |
| ICS algorithm | This paper; Mendeley Data | http://dx.doi.org/10.17632/5hvpvspw82.1 |
| Other | | |
| Sequence data, analyses, and resources related to the ultra-deep sequencing of the AML31 tumor, relapse, and matched normal | This paper | http://aml31.genome.wustl.edu |
| Resource website for the AML31 publication | This paper | https://github.com/chrisamiller/aml31SuppSite |

***PHYSICAL SCIENCE TABLE WITH EXAMPLES FOR AUTHOR REFERENCE***

| **REAGENT or RESOURCE** | **SOURCE** | **IDENTIFIER** |
| --- | --- | --- |
| Chemicals, peptides, and recombinant proteins | | |
| QD605 streptavidin conjugated quantum dot | Thermo Fisher Scientific | Cat#Q10101MP |
| Platinum black | Sigma-Aldrich | Cat#205915 |
| Sodium formate BioUltra, ≥99.0% (NT) | Sigma-Aldrich | Cat#71359 |
| Chloramphenicol | Sigma-Aldrich | Cat#C0378 |
| Carbon dioxide (^13^C, 99%) (<2% ^18^O) | Cambridge Isotope Laboratories | CLM-185-5 |
| Poly(vinylidene fluoride-co-hexafluoropropylene) | Sigma-Aldrich | 427179 |
| PTFE Hydrophilic Membrane Filters, 0.22 μm, 90 mm | Scientificfilters.com/Tisch Scientific | SF13842 |
| Critical commercial assays | | |
| Folic Acid (FA) ELISA kit | Alpha Diagnostic International | Cat# 0365-0B9 |
| TMT10plex Isobaric Label Reagent Set | Thermo Fisher | A37725 |
| Surface Plasmon Resonance CM5 kit | GE Healthcare | Cat#29104988 |
| NanoBRET Target Engagement K-5 kit | Promega | Cat#N2500 |
| Deposited data | | |
| B-RAF RBD (apo) structure | This paper | PDB: 5J17 |
| Structure of compound 5 | This paper; Cambridge Crystallographic Data Center | CCDC: 2016466 |
| Code for constraints-based modeling and analysis of autotrophic *E. coli* | This paper | https://gitlab.com/elad.noor/sloppy/tree/master/rubisco |
| Software and algorithms | | |
| Gaussian09 | Frish et al.^1^ | https://gaussian.com |
| Python version 2.7 | Python Software Foundation | [https://www.python.org](https://www.python.org/) |
| ChemDraw Professional 18.0 | PerkinElmer | <https://www.perkinelmer.com/category/chemdraw> |
| Weighted Maximal Information Component Analysis v0.9 | Rau et al.^2^ | https://github.com/ChristophRau/wMICA |
| Other | | |
| DASGIP MX4/4 Gas Mixing Module for 4 Vessels with a Mass Flow Controller | Eppendorf | Cat#76DGMX44 |
| Agilent 1200 series HPLC | Agilent Technologies | https://www.agilent.com/en/products/liquid-chromatography |
| PHI Quantera II XPS | ULVAC-PHI, Inc. | https://www.ulvac-phi.com/en/products/xps/phi-quantera-ii/ |
